# *FMR1* gene therapy restores activity-driven inhibition and prevents audiogenic seizures in *Fmr1^-/y^* mice

**DOI:** 10.64898/2026.06.12.731901

**Authors:** Beatriz Maio, Aditi Singh, Ralph Hector, Kamal Gadalla, Jim Selfridge, Francesca Aria, Susana R. Louros, Stuart R. Cobb, Emily K. Osterweil

## Abstract

Fragile X syndrome (FXS) is a neurodevelopmental disorder associated with auditory hypersensitivity, circuit hyperexcitability, and seizures. Whether re-expression of the *FMR1* gene and encoded Fragile X Messenger Ribonucleoprotein (FMRP) can restore sensory circuit dysfunction remains unclear. Here, we show that a viral AAV-*FMR1* vector rescues audiogenic seizures in the *Fmr1^⁻/y^* mouse model after both neonatal and adult delivery, indicating that auditory circuit dysfunction remains reversible. Local re-expression in the inferior colliculus (IC) is sufficient to reduce seizure susceptibility, identifying this region as a key site of FMRP-dependent circuit regulation. In the IC, Translating Ribosome Affinity Purification and RNA-seq (TRAP-seq) profiling reveals impaired induction of sound-evoked translation programs in *Fmr1^-/y^* neurons, including those regulated by transcription factor *Npas4*. AAV-*FMR1* restores a WT-like molecular response and normalizes unbalanced sound-evoked activation of VGLUT2+ excitatory neurons over VGAT+ inhibitory neurons in *Fmr1^-/y^* IC. Together, these findings indicate altered translation of *Npas4* in response to sound impairs recruitment of inhibition in the *Fmr1^-/y^* IC, and this can be reversed with AAV-*FMR1* administration. Moreover, the rescue of seizures after adult administration of AAV-*FMR1* supports a gene therapy approach for FXS.

**Highlights:**

- TRAP-seq reveals impaired activity-driven translation in *Fmr1^-/y^* inferior colliculus (IC)
- Deficient *Npas4* induction reduces evoked inhibition in *Fmr1^-/y^* IC
- AAV9-*FMR1* gene therapy normalizes translation and excitatory/inhibitory balance in *Fmr1^-/y^* IC
- AAV9-*FMR1* gene therapy prevents audiogenic seizures in *Fmr1^-/y^* mice when administered neonatally or in adulthood

## Introduction

Fragile X syndrome (FXS) is an X-linked neurodevelopmental disorder and one of the most common monogenic causes of autism spectrum disorder, affecting approximately 1:4000 males and 1:8000 females ^1–6^. Patients with FXS exhibit intellectual disability, sensory hypersensitivity, ADHD, and childhood epilepsy ^5,7^. FXS is caused by CGG repeat expansion in the 5′ UTR of the *FMR1* gene, which leads to transcriptional silencing and loss of the encoded Fragile X Messenger Ribonucleoprotein (FMRP)^2^. FMRP is an RNA-binding protein that negatively regulates mRNA translation ^8–11^, and its loss results in elevated basal protein synthesis in rodent models and patient-derived cells ^12–15^. Although substantial progress has defined the molecular consequences of FMRP loss, how these changes give rise to circuit dysfunction and behavioral phenotypes remains incompletely understood. A major behavioral consequence of hyperactivity in *Fmr1^-/y^* circuits is an increased susceptibility to audiogenic seizures (AGS) in response to elevated sound exposure, which mirrors the enhanced auditory responsiveness found in individuals with FXS ^7,16–22^. Recent studies dissecting the circuit mechanisms contributing to this hyperexcitability show that genetic replacement of *Fmr1* in neurons in the inferior colliculus (IC) is sufficient to reduce susceptibility to AGS in conditional *Fmr1^-/y^* mice ^23^. The IC is a central auditory midbrain hub that integrates sound-related inputs and drives behavioral responses ^24–30^, and it has been implicated in AGS generation in other models ^23,31–34^. Understanding the molecular and cellular mechanisms that go awry in the absence of *Fmr1* in this population is therefore a way to understand sensory hyperexcitability in FXS. Moreover, the reversal of these changes with FMRP re-expression would be good evidence that *FMR1* gene replacement is a viable approach for treating FXS.

In the past decade, gene therapy has shown promise for the treatment of multiple neurodevelopmental disorders and rare diseases, including spinal muscular atrophy, Angelman syndrome, Tuberous Sclerosis, *CDKL5* deficiency disorder, epileptic encephalopathy, and Rett syndrome ^35–43^. These advances have renewed interest in other monogenic neurodevelopmental disorders, especially those that have a clear association between gene expression and clinical outcome. The correlation of FMRP levels with IQ in individuals with FXS suggest that restoring *FMR1* expression could be a successful therapeutic strategy ^44^. However, whether FMRP re-expression can resolve key phenotypes in *Fmr1^-/y^*mice remains unclear. Early studies showing successful re-introduction of the *FMR1* gene in the *Fmr1^-/y^* mouse were complicated by the deleterious effects of FMRP overexpression ^2,45–47^. Subsequent work using AAV9 delivery of human *FMR1* isoforms find that physiological re-expression of FMRP can be achieved, however the impact on key *Fmr1^-/y^* phenotypes is variable ^48–50^. Establishing whether this variability is due to differential efficacy of *FMR1* re-expression is a complex task, because it requires knowledge regarding both the mechanisms by which *FMR1* re-expression is expected to restore function in specific circuits, and an identification of the key circuits that drive behavioral alterations in the *Fmr1^-/y^*mouse.

In this study, we show that AAV9 delivery of a vector containing human *FMR1* (AAV-*FMR1*) is sufficient to restore experience-driven changes in the translatome of IC neurons, and dampens hyperexcitability in IC circuits. Translating Ribosome Affinity Purification and RNA-seq (TRAP-seq) profiling on IC neurons after sound exposure shows the translation profile induced with sound in the WT IC is deficient in the *Fmr1^-/y^* IC, and saturated in the steady-state translatome. There is a notable deficiency in the upregulation of *Npas4*, a transcription factor involved in the strengthening of inhibitory synapses, which is restored by administration of AAV-*FMR1*. Consistent with these findings, interrogation of IC neuron activity after sound exposure reveals an increased activity of VGLUT2+ excitatory versus VGAT+ inhibitory neurons in the *Fmr1^-/y^*IC, which is reversed in mice expressing AAV-*FMR1*. Importantly, AAV-*FMR1* administration significantly reduces the incidence of AGS when delivered neonatally or in adulthood, indicating circuit hyperexcitability in *Fmr1^-/y^* brain is reversible with FMRP replacement later in development. Together, these results identify a novel molecular mechanism driving audiogenic hypersensitivity in the *Fmr1^-/y^* brain and provide support for a gene therapy approach in FXS.

## Results

### AAV-*FMR1* administration results in a stable re-expression of FMRP throughout the *Fmr1^-/y^* brain

To address the feasibility of *FMR1* replacement and to characterize the impact of FMRP re-expression, we developed a single-stranded AAV9 vector (AAV-*FMR1*) encoding the human *FMR1* isoform 7, which it is among the most abundant isoforms in human tissues ^51^. To regulate expression of the *FMR1* transgene, fragments of the human *FMR1* promoter and 3’ untranslated region (3’UTR) were added to facilitate endogenous FMRP expression across different cell and tissue types more accurately than use of a strong, ubiquitous promoter. The 1050 bp fragment of the human *FMR1* promoter has high levels of sequence conservation with other species and is predicted to contain critical regulatory elements needed to drive *FMR1* expression, such as H3K27Ac markers^52^, transcription factor binding motifs^53^ and a CpG island^54^. The 1400 bp fragment of the human *FMR1* 3’UTR is also highly conserved across species and contains putative miRNA recognition sites, which may provide more cell-specific regulation^55^, and the predominant *FMR1* polyadenylation signal. A C-terminal Myc epitope tag was incorporated to facilitate detection of recombinant FMRP (**Fig. 1A**). In our first experiments, we tested the expression of FMRP after delivery of AAV9-*FMR1* versus respective AAV-Null or PBS controls. To test the impact of neonatal delivery, we performed bilateral intracerebroventricular (ICV) injections at postnatal days 0-2 (P0-2) and examined biodistribution and cellular FMRP expression levels at P19–P22 or P58–P61, hereafter referred to as the 3-week and 8-week time points, respectively. In a separate cohort, cellular FMRP expression was assessed 7–10 days after bilateral stereotaxic delivery of AAV-*FMR1* into the IC of adult P50–P53 mice, hereafter referred to as adult stereotaxic IC injections (**Fig. 1B**). Biodistribution of AAV-*FMR1* was analyzed by immunostaining for FMRP and Myc after ICV and stereotaxic IC injections (**Fig. 1C-D**). Our results show that both neonatal and adult delivery of AAV-*FMR1* results in a widespread re-expression of Myc and FMRP across multiple brain regions, including the IC, cortex, and hippocampus, with a neuronal transduction of approximately 60% across genotypes (**Fig. 1E**). These transduction rates are comparable to or greater than those reported in previous gene therapy work ^56^. To quantify the amount of FMRP expressed per cell, we acquired high resolution z-stacks of individual neurons using confocal imaging and quantified signal intensity using IMARIS (**Fig. 1F-H, Fig. S1A**). Our results show FMRP levels in *Fmr1^-/y^* AAV-*FMR1* expressing neurons are < 2-fold greater than WT control neurons in both IC and auditory cortex at the 3-week time point (**Fig. 1F, I**). This expression level remains stable at 8-week time point, with only a slight increase to 2-2.5-fold (**Fig. 1G, I**). Similar expression was observed in the adult stereotaxic IC cohort (**Fig. 1H, I**). Together, these results show that AAV-*FMR1* administration in neonates or adult animals results in a stable physiological expression of FMRP in neurons.

**Figure 1.**
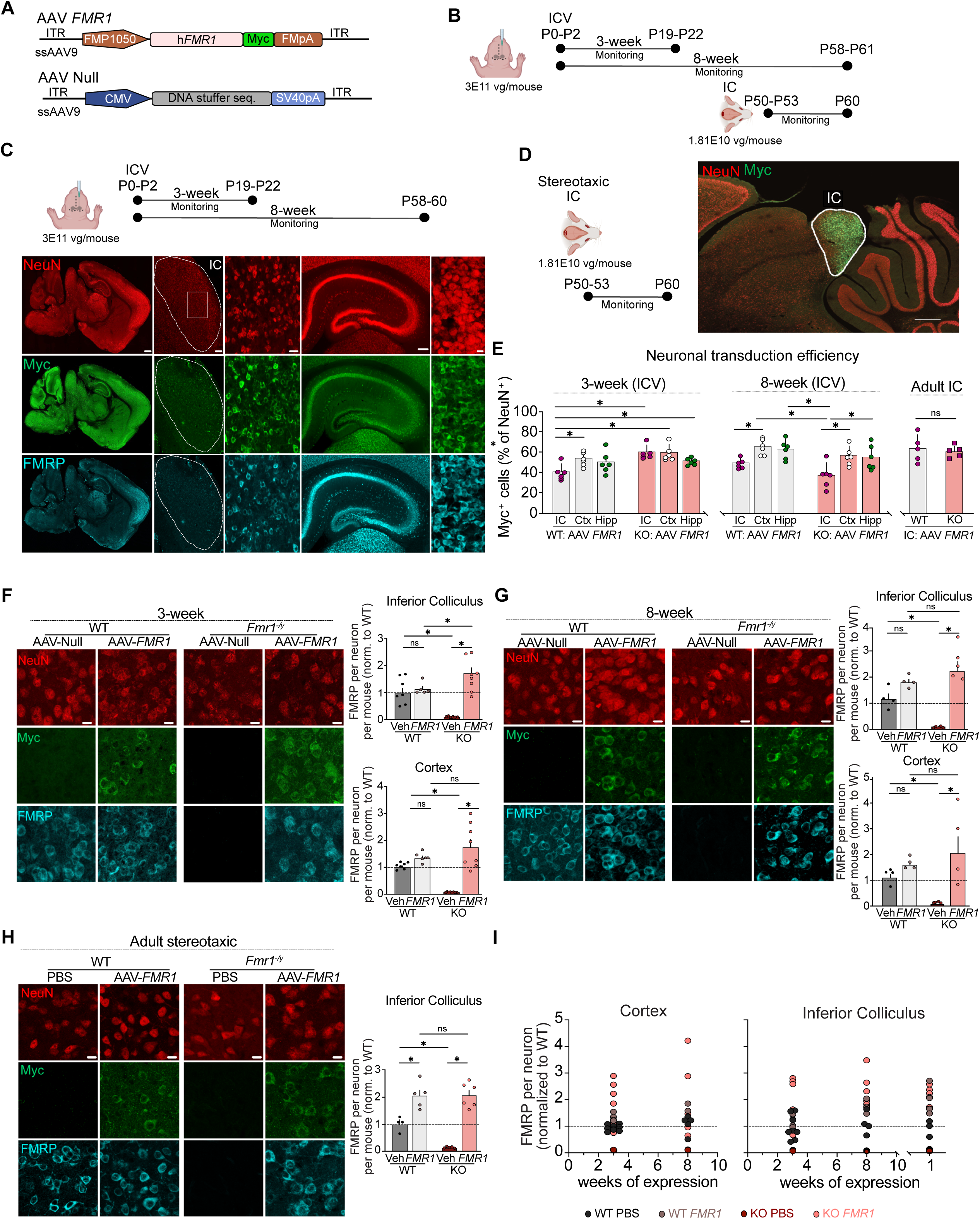
AAV-*FMR1* restores FMRP expression in *Fmr1^⁻/y^* brain. (**A**) Schematic of AAV-*FMR1* and AAV-Null vectors indicates included regulatory components. (**B**) Timelines for FMRP biodistribution testing. Different cohorts of *Fmr1^-/y^* and WT animals were injected with AAV-*FMR1* ICV at P0-P2 and FMRP expression quantified at 3-week and 8-week time points, or injected with AAV-*FMR1* stereotaxically at P50-P53 and FMRP expression quantified after 7-10 days. (**C**) Immunostained sagittal brain section of *Fmr1^-/y^* mouse shows widespread expression of FMRP and AAV-*FMR1* vector (Myc) in neurons (NeuN+) at 8 weeks. Higher magnification images highlight expression in IC and hippocampus. Scale bars left to right: 500 µm, 100 µm, 20 µm, 100 µm and 10 µm. (**D**) Sagittal section of P60 *Fmr1^-/y^* mouse immunostained for Myc shows area of stereotaxic injection. Scale bar = 500 µm. (**E**) AAV-*FMR1* transduction efficiency was quantified as number of Myc+ NeuN+ cells/ total NeuN+ cells. 3 weeks: Two-way ANOVA with Benjamini–Hochberg FDR correction: genotype × region interaction, F(2,30) = 4.77, p = 0.0159; genotype, F(1,30) = 14.54, p = 0.0006; region, F(2,30) = 2.34, p = 0.1140. FDR-significant post hoc comparisons: IC WT versus IC KO, ***p = 0.0006; IC WT versus Ctx WT, *p = 0.0320; IC WT versus Ctx KO, ***q = 0.0006; IC WT versus Hipp KO, *q = 0.0353. 8 weeks: Two-way ANOVA with Benjamini–Hochberg FDR correction: genotype × region interaction, F(2,30) = 0.35, p = 0.7078; region, F(2,30) = 11.54, p = 0.0002; genotype, F(1,30) = 7.59, p = 0.0099. FDR-significant post hoc comparisons: IC WT versus Ctx WT, *p = 0.0172; IC KO versus Ctx WT, ***p = 0.0002; IC KO versus Ctx KO, **p = 0.0079; IC KO versus Hipp WT, **p = 0.0022; IC KO versus Hipp KO, *p = 0.0148. Transduction is similar between acutely injected WT and KO IC, unpaired t-test p = 0.5967. Bars represent mean ± SEM, with each individual point representing the average of 2 sections taken from one mouse. **(F)** Representative 63X images from IC of WT NULL, WT *FMR1*, KO NULL and KO *FMR1* at 3-week time point after ICV injection show expression of Myc and FMRP in neurons of both genotypes. In IC, there was a genotype × treatment interaction, F(1,26) = 20.34, p = 0.0001, and a treatment effect, F(1,26) = 26.41, p < 0.0001, but no genotype effect, F(1,26) = 0.82, p = 0.3725; significant post hoc comparisons were WT PBS versus KO PBS (**p = 0.0010), WT PBS versus KO *FMR1* (**p = 0.0046), KO PBS versus WT *FMR1* (**p = 0.0010), KO PBS versus KO *FMR1* (***p < 0.0001), and WT *FMR1* versus KO *FMR1* (*p = 0.0275). In AuC, there was a genotype × treatment interaction, F(1,25) = 16.38, p = 0.0004, and a treatment effect, F(1,25) = 35.86, p < 0.0001, but no genotype effect, F(1,25) = 2.47, p = 0.1283; significant post hoc comparisons were WT PBS versus KO PBS (***p = 0.0005), WT PBS versus KO *FMR1* (**p = 0.0039), KO PBS versus WT *FMR1* (***p = 0.0001), and KO PBS versus KO *FMR1* (***p < 0.0001). **(G)** Representative 63X images from IC of WT NULL, WT *FMR1*, KO NULL and KO *FMR1* at 8-week time point. In IC, there was a genotype × treatment interaction (p = 0.0131) and a treatment effect (p < 0.0001), but no genotype effect (p = 0.2561); significant post hoc comparisons were WT PBS versus KO PBS (*p = 0.0214), WT PBS versus KO *FMR1* (*p = 0.0205), KO PBS versus WT *FMR1* (**p = 0.0014), and KO PBS versus KO *FMR1* (***p = 0.0001). In AuC, there was a genotype × treatment interaction, F(1,16) = 5.61, p = 0.0308, and a treatment effect, F(1,16) = 13.33, p = 0.0022, but no genotype effect, F(1,16) = 0.96, p = 0.3421; significant post hoc comparisons were WT PBS versus KO PBS (*p = 0.0475), KO PBS versus WT *FMR1* (*p = 0.0153), and KO PBS versus KO *FMR1* (**p = 0.0022). (**H**) Representative 63X images across conditions of IC showing expression of vector and FMRP 7-10 days after stereotaxic injection. Significant interaction (p = 0.0288), together with effects of row factor (p < 0.0001) and column factor (p = 0.0341); post hoc comparisons showed differences between WT PBS and KO PBS (**p = 0.0091), WT PBS and WT *FMR1* (**p = 0.0023), WT PBS and KO *FMR1* (*p = 0.021), KO PBS and WT *FMR1* (***p < 0.0001), and KO PBS and KO *FMR1* (***p < 0.0001), but not between WT *FMR1* and KO *FMR1* (p = 0.9468).All datasets in F-H were analyzed using the same statistical approach. (**I**) FMRP mean intensity across conditions, normalized to WT PBS. Each dot represents the mean of multiple mice. For each mouse, quantification was performed in two sections of the region of interest, with at least 20 neurons analyzed per section, and section values were averaged to obtain a single value per mouse. Data is presented as mean ± SEM. 3 weeks ICV for AuC and IC: WT PBS = 7 mice, WT *FMR1* = 5 mice, KO PBS = 6 mice and KO *FMR1* =8 mice. 8 weeks ICV for AuC and IC: WT PBS = 4 mice, WT *FMR1* = 4 mice, KO PBS = 4 mice and KO *FMR1* = 4 mice. For 1 week in IC: WT PBS = 4 mice, WT *FMR1* = 5 mice, KO PBS = 5 mice and KO *FMR1* = 6 mice.

### Sound-driven translatome changes in WT IC neurons are saturated in the *Fmr1^-/y^* mouse

To understand whether and how the re-expression of FMRP reverses neuropathological changes, we investigated the cellular changes contributing to the increased susceptibility to AGS, one of the most replicable phenotypes in the *Fmr1^-/y^*mouse. Previous work shows that auditory hyperresponsiveness emerges in *Fmr1^-/y^*mice between the second and third postnatal weeks, coinciding with early auditory development and behavioral hypersensitivity ^20,57^. Notably, the incidence of AGS in *Fmr1^-/y^* mice peaks around this time ^13,34,58–61^. While the circuit maturation changes leading to enhanced auditory excitability are likely present in multiple brain areas, it has been shown that genetic replacement of *Fmr1* in *Ntsr1+* neurons of the IC is sufficient to lower the incidence of AGS in *Fmr1^-/y^* mice. We therefore investigated whether *FMR1* re-expression restored cellular alterations in *Fmr1^-/y^* IC neurons using TRAP-seq profiling, an approach that allows for isolation of ribosome-bound mRNAs within specific cell types through conditional expression of an affinity-tagged ribosomal protein^62^. *Fmr1^+/-^*females were bred to a *Snap25*-EGFPL10a line with pan-neuronal expression of GFP-tagged ribosomal L10a. Balanced groups of *Fmr1^-/y^* and WT mice were neonatally injected with AAV-*FMR1* or AAV-Null and TRAP-seq performed on sound-exposed and naïve mice at the 3-week time point (**Fig. 2A**). The sound stimulus consisted of a sampling of mixed frequencies played at >100 dB that we have previously used to evoke AGS in *Fmr1^-/y^*mice ^63^. To mitigate the potential confounds of seizure activity, exposure to the sound stimulus was limited in duration, and stopped short of evoking behavioral seizures. After stimulation, mice were returned to the home cage for 30 min, and IC tissue isolated and processed for TRAP.

**Figure 2.**
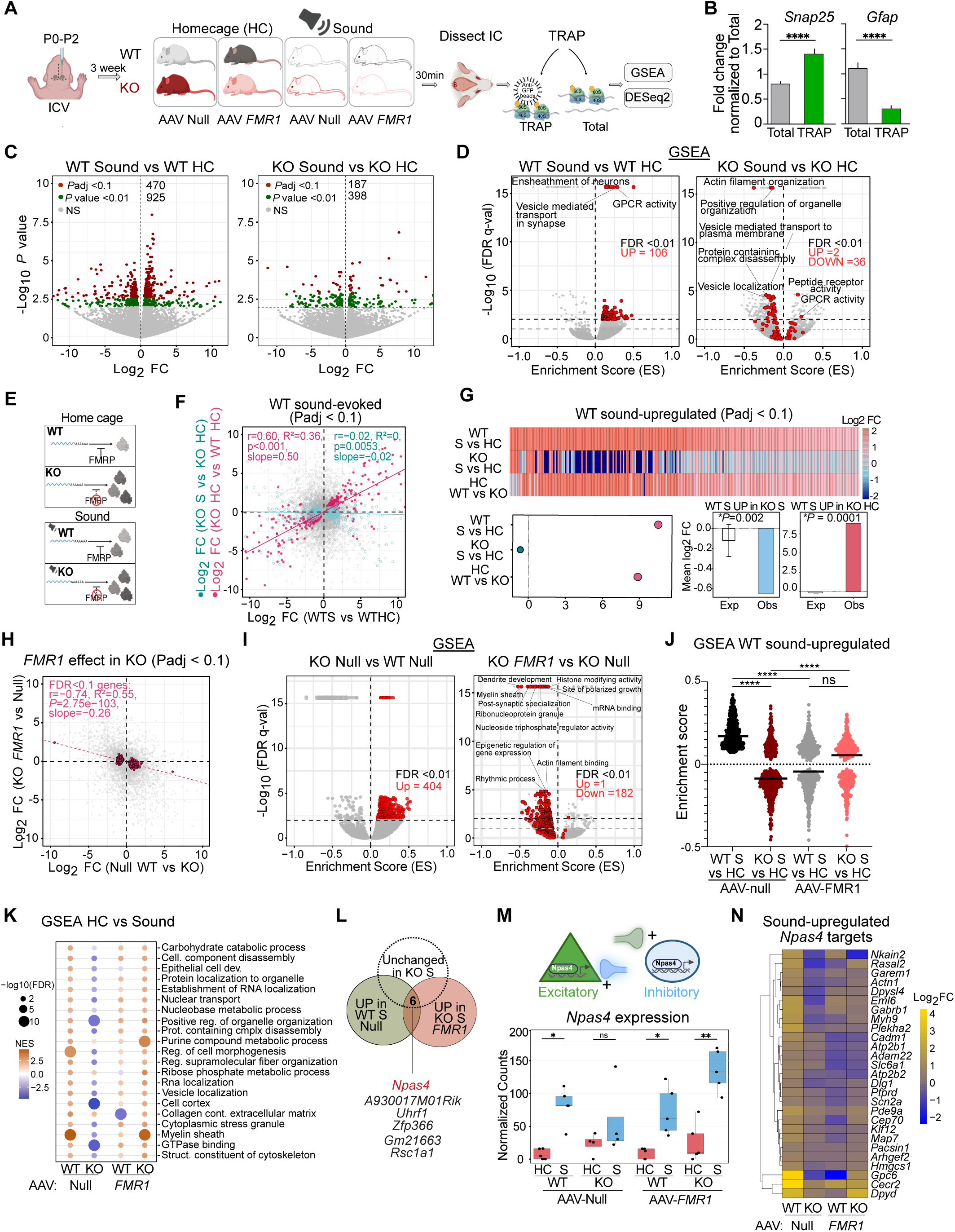
FMRP re-expression restores *Npas4* and activity-driven translation in *Fmr1^⁻/y^* IC neurons. (**A**) Neonatal ICV delivery of AAV9-*FMR1* or AAV9-Null was followed by sound stimulation at 3 weeks, IC collection after 30 min, TRAP, and downstream DESeq2/GSEA analysis. (**B**) qPCR for *Gfap* and *Snap25* indicate neuronal enrichment in TRAP fraction. (**C**) Volcano plots for WT Sound versus HC and KO Sound versus HC comparisons. WT mice show a robust sound-responsive translational program (Padj < 0.1, 470), whereas KO mice show markedly fewer significant changes (Padj < 0.1, 187). (**D**) Volcano plots show gene sets identified by GSEA in sound-evoked WT and sound-evoked KO animals. A comparison shows that the majority of gene sets significantly upregulated in WT Sound (FDR < 0.01, 106 up) are downregulated or unchanged in KO Sound (2 up, 36 down). (**E)** Schematic represents working model in which loss of FMRP elevates basal translation in KO neurons, creating a saturated baseline state that limits further sound-evoked translational changes. (**F)** Scatter plot comparing sound-evoked changes in WT to basal changes in KO reveal a positive correlation (r = 0.60, R2 = 0.36, *p < 0.001), while a comparison to sound-evoked changes in KO reveals no correlation (r = −0.02, R2 = 0). (**G**) Heatmap shows WT sound-upregulated genes (Padj < 0.1) are basally upregulated and non-responsive to sound in KO. Permutation analyses of mean log2 FC confirm the depletion of sound-evoked changes in KO is significant (*p = 0.002), as is the basal upregulation in KO (*p = 0.0001). (**H**) Correlation plot shows that basal changes in KO HC (Padj < 0.1) are reversed with *FMR1* replacement (r = −0.74, R2 = 0.55) **(I)** GSEA volcano plots show pathways significantly upregulated in KO (FDR < 0.01, 404 up) are largely reversed in KO *FMR1* (1 up, 182 down) **(J)** Violin plots show gene sets significantly upregulated with sound in WT (FDR < 0.1), with the same gene sets plotted in sound-evoked KO, WT *FMR1* and KO *FMR1* groups (Kruskal Wallis *p < 0.0001, Dunn’s post hoc WT S vs KO S *p < 0.0001, WT S vs WT *FMR1* S *p < 0.0001, KO S vs KO *FMR1* S *p < 0.0001). (**K)** Dot plot summarizing gene sets that are the most significant (FDR<0.01) and upregulated in (NES > 0) in WT S and show similar upregulation (NES > 0) and significance (FDR<0.01) in KO after *FMR1* replacement. These pathways include catabolic processes, organelle organization, RNA transport, vesicle trafficking, extracellular matrix/collagen pathways, and cytoskeletal organization. (**L)** Venn diagram showing overlap between transcripts upregulated by sound in WT (p<0.01), unchanged in KO S vs KO HC, and upregulated in KO *FMR1* after sound stimulation (p<0.01). Overlapping genes include *Npas4*. (**M**) Top: Schematic depicting *Npas4* as an activity-dependent transcription factor that regulates neuronal activity, synaptic plasticity, and inhibitory synapse development through putative cell-type-specific roles in excitatory and inhibitory neurons; and bottom: Boxplot summary of mean expression of *Npas4* from TRAP-seq, showing reduced *Npas4* levels in KO S and restoration following *FMR1* replacement. WT HC NULL vs. WT Sound NULL, * p = 0.012. KO HC NULL vs. KO Sound NULL, ns, p = 0.486. WT HC *FMR1* vs. WT SOUND *FMR1*, * p = 0.012; KO HC *FMR1* vs. KO SOUND *FMR1*, ** p = 0.008. (**N**) Heatmap of the *Npas4* target genes upregulated by sound in WT (P<0.01) across all groups illustrates impaired induction in KO that is partially restored following *FMR1* re-expression.

To confirm the enrichment of neuron-derived ribosomes in our TRAP samples, we quantified *Snap25* and *Gfap* transcripts in immunoprecipitated and total fractions using qPCR (**Fig. 2B**). Following this confirmation, TRAP samples were subjected to low-input library preparation and Illumina sequencing as in previous work ^64,65^ (**Fig. S2A-B**). To determine whether *Fmr1^-/y^* IC neurons have an abnormal translation response to sound stimulation, we compared TRAP samples from AAV-NULL treated, sound-exposed versus home-cage (HC) WT and *Fmr1^-/y^* animals using DESeq2. Our results show sound stimulation induces a robust translational response in WT mice (470 genes, Padj < 0.1) (**Fig. 2C, Table S1**). In contrast, sound exposure results in a markedly reduced response in *Fmr1^-/y^* mouse (187 genes, Padj < 0.1). To further explore the impact of sound exposure we performed Gene Set Enrichment Analyses (GSEA) to compare changes in groups of transcripts associated with specific cellular pathways. This reveals sound stimulation in WT mice engages programs related to synaptic signaling, membrane trafficking, and neuronal activation, whereas these programs are attenuated or absent in stimulated *Fmr1^-/y^* mice (**Fig. 2D, Table S2**).

Previous studies show that the basal elevation in mRNA translation present in *Fmr1^-/y^* neurons can occlude further stimulated translation downstream of group 1 metabotropic glutamate receptors or TrkB receptors ^14,65^. To investigate whether the diminished change in translation seen with sound exposure in *Fmr1^-/y^* IC neurons is related to a basal saturation, we performed correlation analyses to determine if the changes in sound-evoked transcripts exhibited in WT are similar to constitutive changes in KO (**Fig. 2E**). Comparing the log2 fold-change (FC) of transcripts altered with sound in WT (WT HC vs WT S) reveals a correlation with the changes seen in KO HC vs WT HC, indicating the changes are similar (r = 0.6, R^2^ = 0.36) (**Fig. 2F**). In contrast, these changes are not correlated with those in KO HC vs KO S, indicating they are not evoked (R^2^ = 0). Consistent with these results, A comparison of the mean log2 fold-change of all transcripts significantly upregulated in WT S vs WT HC (Padj < 0.1) shows they are reduced in KO HC vs KO S and similarly upregulated in KO HC vs WT HC (**Fig. 2G**). Permutation analyses show that the depletion of WT S evoked transcripts in the KO S population is significant, as is the increased expression of these targets in the KO HC population. Together, these results show that the WT sound-responsive translational program is elevated at baseline but blunted in its further recruitment in *Fmr1^-/y^* IC.

### FMRP re-expression restores activity-driven changes in *Fmr1^-/y^* IC neurons

To examine whether FMRP re-expression impacts the saturated translation changes observed in *Fmr1^-/y^* IC, we compared changes in AAV-*FMR1* versus AAV-Null expressing groups of WT and *Fmr1^-/y^*animals. A comparison between groups treated with AAV-Null reveals a significant population of differentially expressed transcripts, illustrated by ranked fold-change analysis and GSEA (**Fig. S2C-D, Tables S1-S2**). FMRP re-expression results in significant changes in both *Fmr1^-/y^* (Padj < 0.1, 69) and WT (Padj < 0.1, 24) IC TRAP populations (**Fig. S2E-F**). A correlation of the basal changes seen in KO HC to those changed with *FMR1* re-expression revels a striking negative correlation (r = −0.74, R^2^ = 0.55, P = 2.75E-103) (**Fig. 2H**). The transcripts elevated in the *Fmr1^-/y^* TRAP that are reduced by AAV-*FMR1* expression encode a variety of proteins related to related to protein translation and degradation (*Eif3a*, *Ddx3x*, *Nop58*, *Nedd4*, *Usp9x*), cytoskeletal organization (*Hip1r*) and second messenger signaling pathways (*Akap9*, *Dgkh*) (**Fig. S2G**). GO analysis (**Table S3**) and GSEA further shows that transcripts and gene sets significantly upregulated in KO NULL (Padj < 0.01), including those related to mRNA binding, dendrite development, myelin sheath, actin filament organization, and translation at synapse, are downregulated with *FMR1* replacement (**Fig. 2I, Table S2**). Together, these findings identify a signature of altered translation in *Fmr1^-/y^* IC that is enriched for metabolic and biosynthetic programs, and this is reversed by postnatal re-expression of FMRP.

According to our model, the restoration of basal changes in AAV-*FMR1* treated *Fmr1^-/y^* IC should allow stimulated translation to re-emerge. To answer this remaining question, we first analyzed whether the changes evoked with sound in WT NULL animals remained saturated in the KO with AAV-*FMR1*. DESeq2 comparison shows that sound exposure evoked a significant change in translating mRNAs in both WT and KO IC neurons in AAV-*FMR1* expressing animals (**Fig. S2H-I**). An equivalent sound response is seen in WT (Padj < 0.1 = 63, p < 0.01 = 378) and KO (Padj < 0.1 = 53, p < 0.01 = 349), albeit at a level lower than that seen in WT NULL animals. To determine whether the same basal changes in sound-evoked transcripts are seen in KO with FMRP replacement, we compared significant changes in WT S vs WT HC to changes in KO HC *FMR1* vs WT HC *FMR1* (**Fig. S2J**). This revealed no correlation (R^2^ = 0) despite the positive correlation seen in KO HC NULL versus WT HC NULL (r = 0.61, R^2^ = 0.36, p < 0.01) (**Fig. 2SJ**). This suggests that the sound-evoked changes are among those reversed in the KO with *FMR1* re-expression. To interrogate whether the same pathways upregulated with sound in WT are now upregulated with sound in KO re-expressing *FMR1*, we compared the most significantly upregulated GSEA terms in WT S NULL across conditions (Padj < 0.1) (**Fig. 2J, Table S2**). This comparison shows that the increased population in WT S NULL is not completely restored by

*FMR1* re-expression in the *Fmr1^-/y^* IC. This may be in part due to the general reduced response to sound seen in *FMR1*-treated WT and *Fmr1^-/y^*groups (**Fig. S2H-I, Table S1**). However, a comparison between KO NULL and KO *FMR1* groups indicates a significant difference, with KO *FMR1* shifted towards a WT state. To identify the specific sound-evoked changes that are restored with *FMR1* re-expression in the *Fmr1^-/y^* IC, we compared the most significantly upregulated gene sets in WT S that are also significantly upregulated in KO S *FMR1*. This identifies categories related to metabolic function, RNA localization, cytoskeleton and myelin sheath formation, indicating re-engagement of these key components of the WT activity-responsive program (**Fig. 2K, Table S2**). Together, these results indicate that *FMR1* re-expression in *Fmr1^-/y^* IC leads to a partial restoration of sound-evoked changes in WT IC.

Given the restoration of select sound-evoked pathways in *Fmr1^-/y^*IC with *FMR1* re-expression, we next asked which specific transcripts were restored. To pinpoint the transcripts associated with disrupted function in *Fmr1^-/y^* IC that are restored with *FMR1* re-expression, we overlapped significantly changed transcripts (1) upregulated by sound in WT NULL, (2) unchanged or decreased with sound in KO NULL, and (3) upregulated with sound in KO *FMR1* (**Fig. 2L, Table S1**). This identified six candidates, including the transcription factor *Npas4*, which is among a set of immediate early genes (IEGs) upregulated with neuronal activation^66,67^. *Npas4* is distinguished from other IEGs by its role in regulating the transcription of targets that strengthen inhibitory transmission ^66,68,69^, and impaired induction in the *Fmr1^-/y^*IC would be consistent with a hyperexcitability phenotype. Analysis of normalized TRAP-seq counts confirms that sound-induced *Npas4* expression is robust in WT mice, absent in *Fmr1^-/y^* mice, and restored by FMRP re-expression (**Fig. 2M**). The same impaired induction is not observed for other IEGs, suggesting it is not a marker of deficient activity-driven transcription (**Fig. S2K**). To determine whether the expression of *Npas4* targets followed the alterations in *Npas4*, we investigated whether the targets significantly upregulated in WT were also upregulated in other groups (**Fig. 2N**). A heatmap of these changes shows that the majority of *Npas4* targets significantly upregulated in WT exhibit a diminished recruitment in KO that is restored with *FMR1* re-expression (**Fig. 2N**). The GO terms represented by these targets are largely related to synaptic function (**Fig. S2K, Table S4)**. Together, these results identify *Npas4*, a regulator of synaptic inhibition, as key element of the sound-evoked translation profile in WT IC neurons that is impaired in *Fmr1^-/y^* and restored with FMRP re-expression.

### Impaired induction of *Npas4* in *Fmr1^-/y^* IC neurons is restored with FMRP re-expression

Our TRAP-seq results highlight an impaired upregulation of *Npas4* in *Fmr1^-/y^* IC neurons that is restored to WT levels with AAV-*FMR1*. To validate this effect, we quantified *Npas4* expression in the IC of additional groups of WT and *Fmr1^-/y^* IC mice using RNAscope and probes to *Npas4* (**Fig. 3A-B**). Although this method quantifies changes in RNA expression rather than ribosome engagement, *Npas4* transcript is known to increase dramatically in both transcription and translation, both of which are likely read-out as a change in the TRAP fraction. WT and *Fmr1^-/y^* littermates were exposed to sound as for TRAP-seq experiments, then returned to the home cage for 45 min and perfuse-fixed brains isolated and sectioned. Analyses of confocal z-stacks using IMARIS reveal a significant increase in the density of *Npas4*+ neurons in WT IC after exposure to sound (**Fig. 3C, Fig. S3A**). In *Fmr1^-/y^*IC, there is a basal elevation in the density of *Npas4*^+^ neurons, with no further increase seen after sound stimulation. These results are consistent with TRAP-seq results showing induction of *Npas4* in WT that is impaired in *Fmr1^-/y^*. Previous studies of *Npas4* show that it activates transcription profiles to strengthen inhibitory synapses in both glutamatergic excitatory and GABAergic inhibitory neurons^66^. As our TRAP-seq experiments were performed on a combined population, we wondered if the impaired recruitment of *Npas4* resided specifically in glutamatergic or GABAergic populations. To assess this, we quantified the expression of *Npas4* in VGLUT2 and VGAT neuron subtypes after sound stimulation using RNAscope. VGLUT2+ neurons represent the greatest population of excitatory neurons in the IC, and they are reciprocally connected to VGAT+ neurons that inhibit activity both locally and through long-range projections ^70^. Our quantification reveals a higher expression of *Npas4* within VGAT+ populations (∼60% cells) as compared to VGLUT2⁺ populations (∼10% cells) in the IC (**Fig. 3D-E**). Nevertheless, the recruitment of *Npas4* with sound is impaired in both VGLUT2^+^ and VGAT^+^ populations of the *Fmr1^-/y^* IC, with basal upregulation in both cell types (**Fig. 3D-E)**. This indicates the impairment in *Npas4*-mediated transcriptional programs is present in both glutamatergic and GABAergic cell types in the *Fmr1^-/y^* IC.

**Figure 3.**
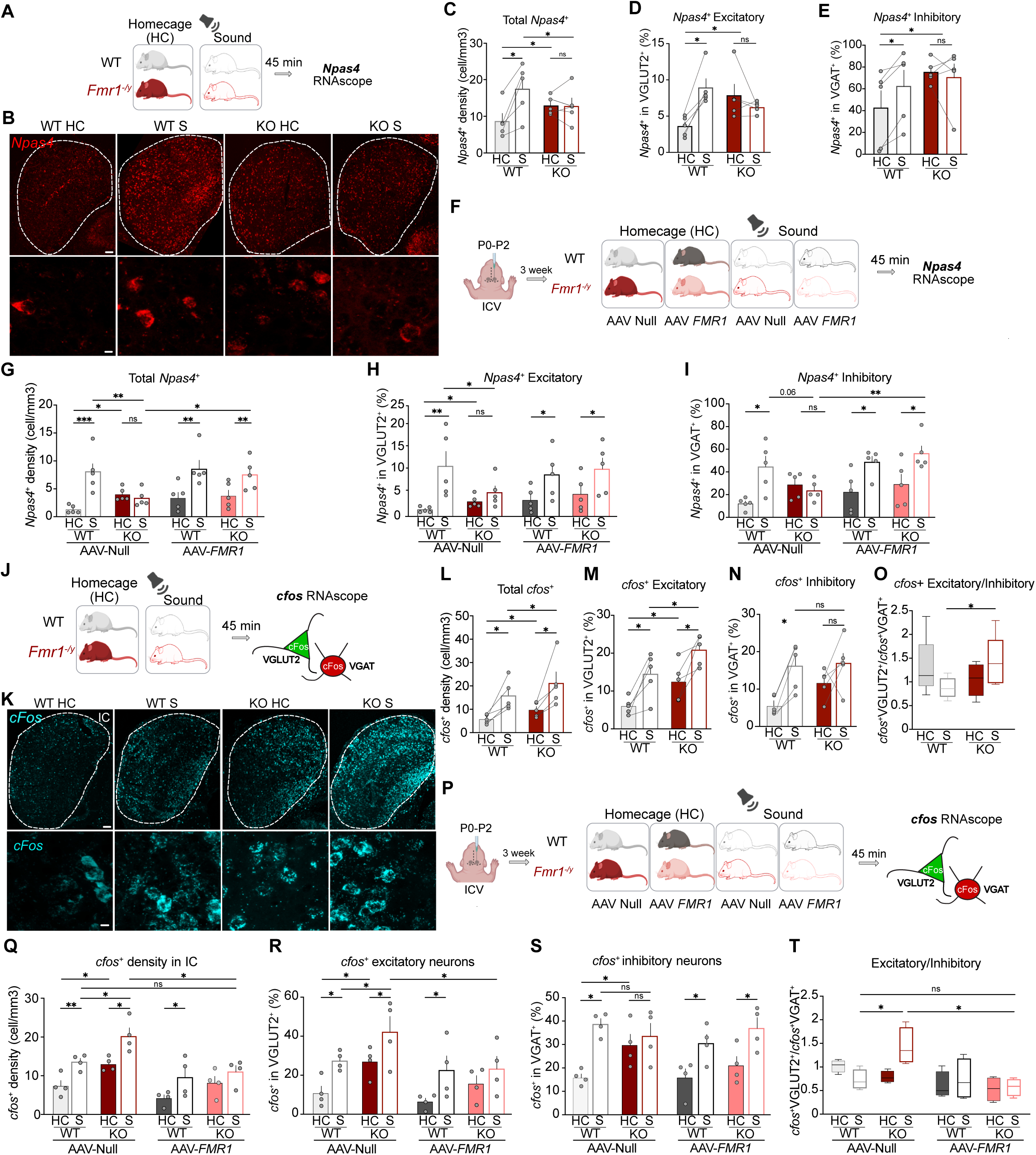
FMRP replacement rescues sound-evoked excitatory/inhibitory neuronal activation balance in the *Fmr1^⁻/y^* IC. (**A**) Untreated 3-week-old WT and *Fmr1^⁻/y^* mice were exposed to sound or kept in home cage and perfuse-fixed 45 min later. IC sections were processed for RNAscope analysis for *Npas4.* (**B**) Representative IC sections across genotypes and conditions, probed for *Npas4*, VGLUT2 and VGAT. Higher-magnification views show VGLUT2^+^, VGAT^+^, and *Npas4* ^+^. Scale bar from top to bottom, 100um and 10um. (**C**) Quantification of *Npas4*⁺ cell density in the IC in WT and *Fmr1^⁻/y^* mice following sound or home cage condition. WT HC, 7.73 ± 2.48; WT Sound, 15.94 ± 3.00; KO HC, 11.18 ± 2.23; KO Sound, 12.90 ± 2.18. Two-way ANOVA: genotype × stimulus interaction, F(1,4) = 30.47, p = 0.0053. Post hoc comparisons: WT HC vs. WT Sound, **p = 0.0035; WT HC vs. KO HC, *p = 0.0214; WT Sound vs. KO Sound, *p = 0.0260; KO HC vs. KO Sound, ns, p = 0.1065. (**D**) Quantification of the percentage of VGLUT2⁺ neurons that were *Npas4*⁺ in the IC from WT and *Fmr1^⁻/y^* across conditions. WT HC, 3.60 ± 0.61; WT Sound, 8.93 ± 1.23; KO HC, 7.86 ± 1.58; KO Sound, 6.20 ± 0.34. Two-way ANOVA: genotype × stimulus interaction, F(1,4) = 23.39, p = 0.0084. Post hoc comparisons: WT HC vs. WT Sound, *p = 0.0387; WT HC vs. KO HC, *p = 0.0422; KO HC vs. KO Sound, ns, p = 0.2157. (**E**) Quantification of the percentage of VGAT⁺ neurons that were *Npas4*⁺ in the IC from WT and *Fmr1^⁻/y^* across conditions. WT HC, 42.58 ± 15.85; WT Sound, 62.42 ± 14.94; KO HC, 75.38 ± 4.88; KO Sound, 70.51 ± 12.71. Two-way ANOVA: genotype × stimulus interaction, F(1,4) = 9.811, p = 0.0351. Post hoc comparisons: WT HC vs. WT Sound, *p = 0.0473; WT HC vs. KO HC, *p = 0.0223; KO HC vs. KO Sound, ns, p = 0.4321. (**F**) P21 WT and *Fmr1^⁻/y^* mice, treated at P0-2 with AAV Null or AAV-*FMR1* were exposed to sound or kept in home cage, perfused 45 min later. (**G**) Quantification of *Npas4*⁺ cell density in the IC in WT and *Fmr1^⁻/y^* mice across conditions. WT HC NULL, 1.33 ± 0.32; WT Sound NULL, 8.10 ± 1.64; WT HC *FMR1*, 3.47 ± 1.11; WT Sound *FMR1*, 8.56 ± 1.55; KO HC NULL, 3.76 ± 0.67; KO Sound NULL, 3.33 ± 0.67; KO HC *FMR1*, 3.76 ± 1.02; KO Sound *FMR1*, 7.53 ± 1.24. Three-way ANOVA: stimulation, F(1,8) = 39.97, p = 0.0002; genotype × stimulation, F(1,8) = 18.63, p = 0.0026; genotype × treatment × stimulation, F(1,8) = 8.875, p = 0.0176. Post hoc comparisons: WT HC NULL vs. WT Sound NULL, ***p = 0.0004; WT HC NULL vs. KO HC NULL, *p = 0.0490; WT Sound NULL vs. KO Sound NULL, **p = 0.0016; WT HC *FMR1* vs. WT Sound *FMR1*, **p = 0.0016; KO HC NULL vs. KO Sound NULL, ns, p = 0.8806; KO Sound NULL vs. KO Sound *FMR1*, *p = 0.0238; KO HC *FMR1* vs. KO Sound *FMR1*, **p = 0.0097. (**H**) Quantification of the percentage of VGLUT2⁺ neurons that were *Npas4*⁺ in the IC. WT HC NULL, 1.55 ± 0.28; WT Sound NULL, 11.35 ± 3.29; WT HC *FMR1*, 3.70 ± 1.24; WT Sound *FMR1*, 9.41 ± 2.39; KO HC NULL, 3.38 ± 0.61; KO Sound NULL, 5.41 ± 1.47; KO HC *FMR1*, 5.07 ± 1.78; KO Sound *FMR1*, 10.67 ± 2.48. Three-way ANOVA: stimulation, F(1,8) = 19.89, p = 0.0021; genotype × treatment, F(1,8) = 7.915, p = 0.0227; genotype × stimulation, F(1,8) = 7.406, p = 0.0262; genotype × treatment × stimulation, F(1,8) = 7.027, p = 0.0292. Post hoc comparisons: WT HC NULL vs. WT Sound NULL, **p = 0.0073; WT HC NULL vs. KO HC NULL, *p = 0.0490; WT Sound NULL vs. KO Sound NULL, **p = 0.0016; WT HC *FMR1* vs. WT Sound *FMR1*, *p = 0.0470; KO HC NULL vs. KO Sound NULL, ns, p = 0.4867; KO HC *FMR1* vs. KO Sound *FMR1*, *p = 0.0470. (**I**) Quantification of the percentage of VGAT⁺ neurons that were *Npas4*⁺ in the IC. WT HC NULL, 12.15 ± 2.24; WT Sound NULL, 44.59 ± 9.34; WT HC *FMR1*, 22.34 ± 7.90; WT Sound *FMR1*, 48.73 ± 5.07; KO HC NULL, 28.91 ± 5.14; KO Sound NULL, 23.58 ± 4.00; KO HC *FMR1*, 29.23 ± 8.69; KO Sound *FMR1*, 56.25 ± 6.62. Three-way ANOVA: stimulation, F(1,8) = 30.89, p = 0.0005. Post hoc comparisons: WT HC NULL vs. WT Sound NULL, *p = 0.0233; WT HC *FMR1* vs. WT Sound *FMR1*, *p = 0.0378; KO HC NULL vs. KO Sound NULL, ns, p = 0.6109; KO Sound NULL vs. KO Sound *FMR1*, **p = 0.0089; KO HC *FMR1* vs. KO Sound *FMR1*, *p = 0.0378.n = 5 littermate pairs. Data are presented as mean ± SEM. (**J**) P21 WT and *Fmr1^⁻/y^*mice were exposed to sound stimulation or maintained under home-cage conditions and perfused 45 min later probed for *cfos.* (**K**) Representative IC sections across genotypes and conditions. Higher-magnification views show *cfos^+^*. Scale bar from top to bottom, 100 µm and 10 µm. (**L**) *cfos*⁺ hyperactivation of *Fmr1^⁻/y^* upon sound shown by cell density in the IC. WT HC, 5.72 ± 0.86; WT S, 15.61 ± 2.98; KO HC, 10.10 ± 1.19; KO S, 21.41 ± 4.54. Two-way ANOVA: genotype, F(1,4) = 8.087, p = 0.0467; stimulus, F(1,4) = 10.63, p = 0.0311. Post hoc comparisons: WT HC vs. WT S, **p = 0.0011; WT HC vs. KO HC, *p = 0.0108; WT S vs. KO S, **p = 0.0058; KO HC vs. KO S, *p = 0.0009. (**M**) Quantification of the percentage of VGLUT2⁺ neurons that were cfos⁺ in the IC hyperactivation of *Fmr1^⁻/y^* upon sound. WT HC, 6.04 ± 0.98; WT S, 14.49 ± 2.60; KO HC, 12.13 ± 2.02; KO S, 20.86 ± 1.80. Two-way ANOVA: stimulus, F(1,4) = 15.3, p = 0.0173. Post hoc comparisons: WT HC vs. WT S, **p = 0.0010; WT HC vs. KO HC, **p = 0.0021; WT S vs. KO S, **p = 0.0021; KO HC vs. KO S, **p = 0.0010. (**N**) Quantification of the percentage of VGAT⁺ neurons that were cfos⁺ in the IC. WT HC, 5.33 ± 1.28; WT S, 16.53 ± 2.86; KO HC, 12.26 ± 1.79; KO S, 16.50 ± 2.98. Two-way ANOVA: stimulus, F(1,4) = 8.764, p = 0.0415. Post hoc comparisons: WT HC vs. WT S, *p = 0.0173; KO HC vs. KO S, ns, p = 0.1329; WT S vs. KO S, ns, p = 0.9888. Each dot represents one mouse, with values averaged across two sections per animal. (**K-N**) n = 5 littermate pairs. Data are presented as mean ± SEM. (**O**) Excitation/Inhibition (E/I) balance increased in *Fmr1^⁻/y^* upon sound. Mann Whitney test, * p = 0.0317. (**P**) P21 WT and *Fmr1^⁻/y^*mice, at P0–P2 ICV-injected, were exposed to sound or kept in home cage, perfused 45 min later. (**Q**) Quantification of cfos⁺ cell density in the IC in WT and *Fmr1^⁻/y^* mice following AAV Null or AAV *FMR1* treatment. WT NULL HC, 7.33 ± 1.42; WT NULL S, 13.52 ± 0.97; WT *FMR1* HC, 4.16 ± 0.92; WT *FMR1* S, 9.53 ± 2.51; KO NULL HC, 12.95 ± 0.90; KO NULL S, 20.14 ± 2.11; KO *FMR1* HC, 8.13 ± 1.71; KO *FMR1* S, 11.00 ± 1.65. Three-way ANOVA: genotype, F(1,6) = 14.39, p = 0.0090; treatment, F(1,6) = 10.49, p = 0.0177; stimulus, F(1,6) = 98.25, p < 0.0001. Post hoc comparisons: WT NULL HC vs. WT NULL S, *p = 0.0211; WT NULL HC vs. KO NULL HC, *p = 0.0476; KO NULL HC vs. KO NULL S, *p = 0.0116; WT NULL S vs. KO NULL S, *p = 0.0237; WT *FMR1* HC vs. WT *FMR1* S, *p = 0.0328; KO NULL S vs. KO *FMR1* S, **p = 0.0025. (**R**) Quantification of the percentage of VGLUT2⁺ neurons that were cfos⁺ in the IC. WT NULL HC, 14.08 ± 2.76; WT NULL S, 27.34 ± 2.38; WT *FMR1* HC, 7.28 ± 1.12; WT *FMR1* S, 22.51 ± 7.40; KO NULL HC, 26.88 ± 3.82; KO NULL S, 42.12 ± 8.06; KO *FMR1* HC, 15.60 ± 4.21; KO *FMR1* S, 23.23 ± 6.48. Three-way ANOVA: genotype, F(1,6) = 11.41, p = 0.0149; stimulus, F(1,6) = 29.94, p = 0.0016. Post hoc comparisons: WT NULL HC vs. WT NULL S, *p = 0.0321; WT NULL HC vs. KO NULL HC, *p = 0.0497; KO NULL HC vs. KO NULL S, *p = 0.0262; WT NULL S vs. KO NULL S, *p = 0.0321; WT *FMR1* HC vs. WT *FMR1* S, *p = 0.0262; KO NULL S vs. KO *FMR1* S, *p = 0.0353. (**S**) Quantification of the percentage of VGAT⁺ neurons that were cfos⁺ in the IC. WT NULL HC, 15.51 ± 1.45; WT NULL S, 38.63 ± 2.44; WT *FMR1* HC, 15.81 ± 3.84; WT *FMR1* S, 30.43 ± 3.89; KO NULL HC, 29.83 ± 4.62; KO NULL S, 33.54 ± 5.51; KO *FMR1* HC, 19.99 ± 3.03; KO *FMR1* S, 36.92 ± 4.58. Three-way ANOVA: stimulus, F(1,12) = 28.26, p = 0.0002. Post hoc comparisons: WT NULL HC vs. WT NULL S, **p = 0.0048; WT NULL HC vs. KO NULL HC, *p = 0.0480; KO NULL HC vs. KO NULL S, ns, p = 0.6133; WT NULL S vs. KO NULL S, ns, p = 0.5174; WT *FMR1* HC vs. KO *FMR1* HC, ns, p = 0.5831; WT *FMR1* HC vs. WT *FMR1* S, *p = 0.0375; KO *FMR1* HC vs. KO *FMR1* S, *p = 0.0174. Each dot represents one mouse, with values averaged across two sections per animal. n = 4 littermate pairs. Data are presented as mean ± SEM. (**T**) Excitation/Inhibition (E/I) balance of WT and *Fmr1^⁻/y^*across conditions. Mann Whitney test, WT NULL S vs. KO NULL S, * p = 0.0286; KO NULL S vs. KO *FMR1* S, * p = 0.0286. Data are presented as mean ± SEM.

Our TRAP-seq data show that impaired recruitment of *Npas4* in *Fmr1^-/y^* IC neurons is restored by FMRP re-expression. To validate this, we further examined *Npas4*^+^ neuron density in WT and *Fmr1^-/y^* mice after neonatal injection of AAV-Null or AAV-*FMR1* (**Fig. 3F)**. This reveals the impaired recruitment of *Npas4*+ neurons in *Fmr1^-/y^* IC is restored to WT levels at the 3-week timepoint re-expression of FMRP (**Fig. 3G).** The restored induction of *Npas4* is seen for both VGLUT2+ and VGAT+ populations (**Fig. 3H-I)**. Together, our results confirm that impaired induction of *Npas4* in the *Fmr1^-/y^* IC after sound exposure is normalized with AAV-*FMR1*.

### Replacement of FMRP restores imbalanced activation of VGLUT2+ versus VGAT+ neurons in the *Fmr1^-/y^* IC in response to sound

Although we and others find an increased activation of IC neurons in the *Fmr1^-/y^* mouse in response to sound stimulation ^71^, the mechanisms linking this change to AGS are not well understood. Previous studies in epilepsy prone rats and mice indicate local circuit activation within the IC is particularly relevant for AGS generation and the motor outputs associated with behavioral expression^72–74^. The upregulation of *Npas4* upon sound stimulation in WT IC neurons suggests a recruitment of inhibition that is impaired in the *Fmr1^-/y^* IC. We therefore investigated whether there was an imbalance in the activation of VGLUT2+ excitatory and VGAT+ inhibitory neurons in the *Fmr1^-/y^* IC. The activity driven response was assessed using in situ profiling with RNAscope probes to *cfos*. Similar to our analysis of *Npas4* activation, balanced groups of *Fmr1^-/y^* and WT mice were exposed to sound and returned to the home cage for 45 min (**Fig. 3J**). The density of *cfos+* neurons was compared between sound-exposed and home cage animals similar to previous work^63^. Our results show that sound stimulation significantly increases the density of *cfos+* neurons in the IC, specifically within the core region that contains ascending projections to the auditory cortex (**Fig. 3J**)^75^. Quantification of *cfos+* neuron density reveals a significantly greater response in *Fmr1^-/y^* versus WT IC (**Fig. 3K-L, Fig. S3A**). To determine whether this effect is seen in excitatory neurons, we quantified the proportion of *cfos+* within the VGLUT2+ population of the IC. Our results show that sound stimulation increases activation of VGLUT2+ neurons in both genotypes, with a larger total activation level observed in the *Fmr1^-/y^* IC (**Fig. 3M-N**). Additionally, basal activation of VGLUT2+ neurons is higher in *Fmr1^-/y^* IC. In contrast, while an increase in the activation in VGAT+ neurons is seen in WT, this difference fails to reach significance in *Fmr1^-/y^* IC (**Fig. 3M-N**). No difference is seen in total numbers of VGLUT2+ or VGAT+ neurons in the IC across genotypes (**Fig. S3 E-F**).

To determine the differential activation of VGLUT2+ versus VGAT+ populations suggestive of local excitatory/inhibitory (E/I) balance, we compared the ratio of cfos+ VGLUT2+: VGAT+ neurons in HC WT versus *Fmr1^-/y^* IC. Analysis of this ratio in HC animals did not reveal a significant difference, suggesting that although there is a basal increase in the activation of both populations in WT versus *Fmr1^-/y^*, these changes are balanced with one another. However, sound stimulation induces a significant increase in VGLUT2+: VGAT+ activation in *Fmr1^-/y^* IC versus WT, indicating a net increase in excitatory versus inhibitory activity (**Fig. 3O, Fig. S3G-H)**. Together, these results suggest impaired activation of VGAT+ neurons results in elevated E/I activation in *Fmr1^-/y^* IC in response to sound, consistent with the reduced recruitment of *Npas4* seen in our TRAP-seq profiling.

As the impaired induction of *Npas4* is reversed with AAV-*FMR1*, we wondered if this treatment would similarly restore inhibitory activation and E/I balance in the *Fmr1^-/y^* IC. We examined the activation of IC neurons after sound stimulation in an independent cohort of WT and *Fmr1^-/y^* animals neonatally injected with AAV-*FMR1* or AAV-Null as done for *Npas4* RNAscope experiments (**Fig. 3P**). Our results show that, similar to uninjected animals, AAV-Null expressing *Fmr1^-/y^*animals exhibit a significant increase in sound-evoked IC neuron activation versus WT animals (**Fig. 3Q**). In AAV-*FMR1* expressing *Fmr1^-/y^* animals, this level of activation is significantly reduced to WT levels. Quantification of VGLUT2^+^ neurons shows a similar increase in stimulated activation in AAV-Null treated *Fmr1^-/y^* IC that is significantly reduced in AAV-*FMR1* treated *Fmr1^-/y^* mice (**Fig. 3R**). In contrast to the VGLUT2+ population, sound stimulation does not significant increase activation of VGAT^+^ neurons in AAV-Null treated *Fmr1^-/y^*animals, whereas AAV-*FMR1* treatment results restores increased VGAT+ activation to WT levels (**Fig. 3S**). A comparison of VGLUT2+: VGAT+ activation across conditions shows that AAV-*FMR1* treatment reduces the activity-driven elevation in E/I seen in *Fmr1^-/y^* back down to WT levels (**Fig. 3T, Fig. S3G-H**). These results suggest that re-expression of FMRP enables the recruitment of VGAT+ neurons in response to sound stimulation in *Fmr1^-/y^* IC, restoring the E/I balance to WT levels.

### *FMR1* replacement suppresses AGS when delivered early or late in development

Having established that AAV-*FMR1* restores impaired *Npas4* expression and recruitment of VGAT+ inhibitory neurons upon sound stimulation, we next asked whether this treatment prevents AGS (**Fig. 4A**). Neonatal ICV injections were performed on *Fmr1^-/y^*and WT groups and AGS tested at 3-week or 8-week time point, as in previous studies ^64,71,76^. Animals were transferred to a test chamber where they were habituated for 1 min, then exposed to a 2-min sampling of a defined mixture of frequencies played at >100 dB. Seizures were scored on a 1-3 scale of increasing severity: (1) wild running (WR; pronounced, undirected running and thrashing), (2) clonic seizure (violent spasms accompanied by loss of balance), and (3) tonic seizure (postural rigidity in limbs) (**Fig. 4B; Videos S1-S16).** Consistent with previous studies, we find that PBS-treated *Fmr1^-/y^* mice exhibit a greater incidence of AGS relative to WT littermates at both 3 and 8-week time point (**Fig. 4C–E**). Neonatal AAV9-*FMR1* delivery significantly reduces AGS incidence at both ages, whereas AAV-Null shows comparable seizure incidence to PBS controls (**Fig. 4C-D**). This protective effect can be seen in the most severe seizure category, with tonic seizures reduced from 91.7% to 15.4% at 3 weeks and from 61.1% to 7.7% at 8 weeks. These results show that neonatal *FMR1* re-expression lowers AGS susceptibility in *Fmr1^-/y^* mice at both juvenile and adult ages.

**Fig 4.**
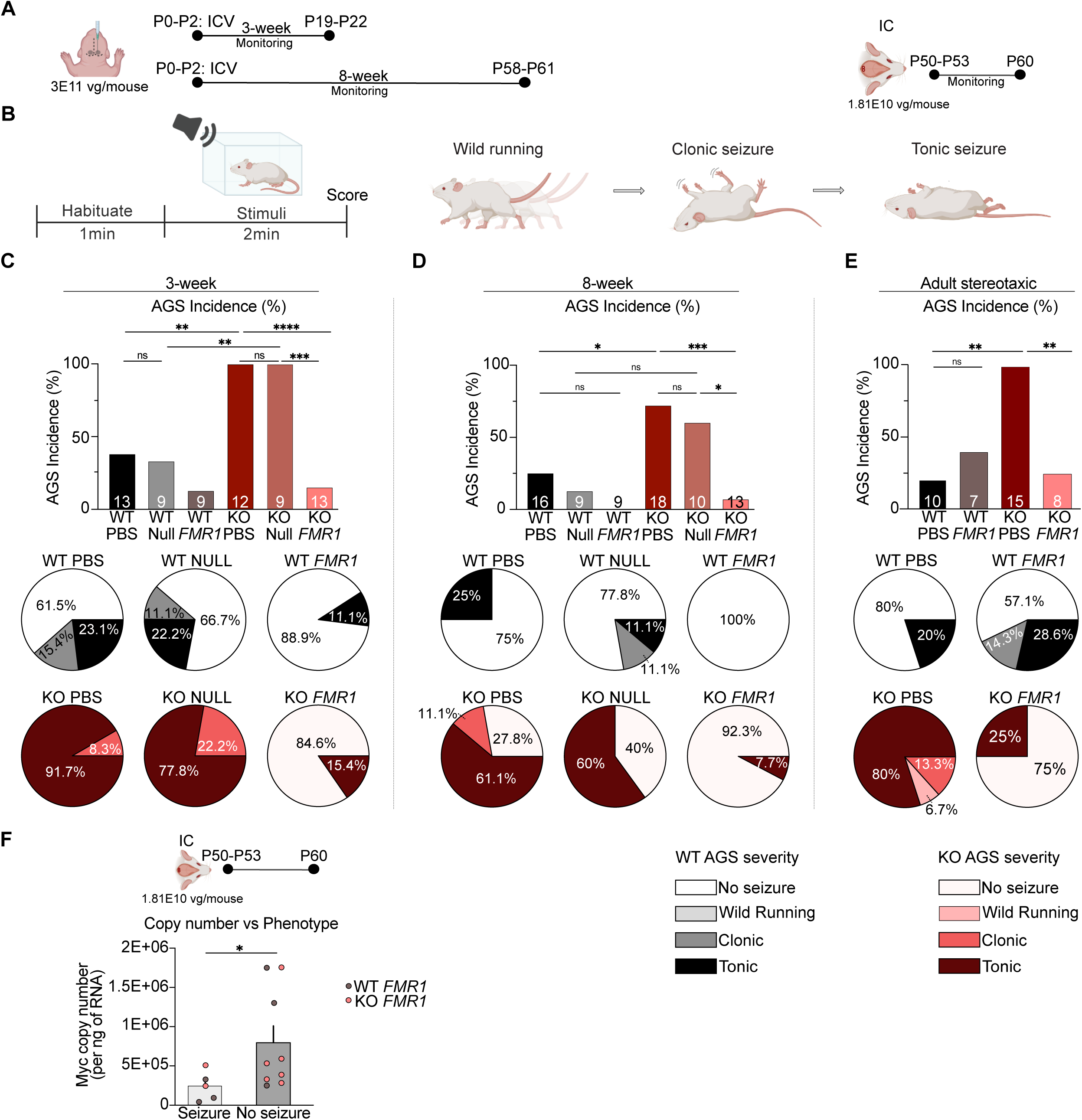
FMR1 replacement suppresses AGS following acute IC delivery and short- or long-term ICV treatment. (**A**) Experimental design of AAV-*FMR1* paradigms, including neonatal ICV delivery at P0–P2 with 3- or 8-week time point expression and adult local IC delivery at P50-P53 with 7-10 days of expression. (**B**) Schematic of AGS assay, showing habituation of mice for 1min followed by stimuli during 2min. Increasing severity was score as wild running, clonic seizure and tonic seizure. (**C**) AAV-*FMR1* after 3 weeks expression reduces the incidence of AGS in *Fmr1^-/y^*mice. AAV-Null and PBS ICV showing comparable AGS incidence. Fisher’s exact test, WT PBS vs. KO PBS ∗∗p=0.0016, KO PBS vs. KO *FMR1*∗∗∗∗p<0.0001, KO Null vs. KO *FMR1*∗∗∗p=0.0002, WT Null vs. KO Null ∗∗p=0.0014). (**D**) AAV-*FMR1* reduces the incidence of AGS in *Fmr1^-/y^* mice. Fisher’s exact test, WT PBS vs. KO PBS ∗p=0.0149, KO PBS vs. KO *FMR1*∗∗∗p=0.0008, KO Null vs. KO *FMR1*∗∗p=0.0088). (**E**) Adult stereotaxic IC expression of AAV-*FMR1* reduces the incidence of AGS in *Fmr1^-/y^* mice at p60. Fisher’s exact test, WT PBS vs KO PBS **p=0.0043. KO PBS vs KO *FMR1* **p=0.0017. Number in each bar indicate the number of mice tested per condition. Pie charts show the distribution of behavioral outcomes within the indicated groups, expressed as the percentage of the total number of mice tested. AAV-*FMR1* reduces tonic seizures in *Fmr1^-/y^* mice. (**F**) Myc copy number in the IC of animals following stereotaxic AAV *FMR1* delivery reveals that those with no seizures have a significantly higher Myc copy number versus those that exhibit seizures (*p =0.0420). Data are presented as mean ± SEM; each dot represents one mouse (n = 5 seizure, n = 9 no seizure).

An important question regarding gene therapy is the extent to which deleterious symptoms can be reversed with a treatment delivered later in development. We therefore tested whether re-expression of FMRP in adult animals could prevent AGS by performing stereotaxic injections of AAV-*FMR1* or PBS in *Fmr1^-/y^* and WT mice at P50-P53 and test AGS 7-10 days after injection. Our results show that PBS-treated *Fmr1^-/y^*mice exhibit a seizure incidence of 100% with 80% exhibiting tonic seizures (**Fig. 4E**). In contrast, only 25% of AAV-*FMR1* expressing mice exhibit seizures when tested 7-10 days after injection. Additionally, qPCR analysis of IC tissue taken from animals after AGS testing reveals a significant elevation in Myc expression in animals that did not exhibit seizures versus those that did, indicating the level of FMRP re-expression is correlated with AGS outcome (**Fig. 4F**). Together, these findings re-affirm the critical role of FMRP expression in the IC for preventing AGS and show that AAV-*FMR1* administration in adult animals normalizes a central hyperexcitability phenotype in *Fmr1^-/y^* mice.

## Discussion

FXS is widely associated with circuit hyperexcitability, but the mechanisms linking loss of FMRP to sensory-driven dysfunction remain incompletely understood. Our findings support a model in which FMRP enables auditory input to engage adaptive molecular responses in the IC that recruits *Npas4* to promote circuit stability. In *Fmr1^-/y^* mice, this coupling is disrupted, leading to blunted sound-evoked translation, enhanced stimulus-induced excitation due to impaired inhibitory neuron activation, and increased expression of AGS. Re-expression of FMRP normalizes these alterations, indicating substantial reversibility of IC circuit dysfunction. Together with prior work implicating the IC in AGS susceptibility, these results identify restoration of FMRP function in the IC as a mechanism for normalizing sound-evoked excitatory/inhibitory balance and activity-dependent gene regulation ^23,71,75^. Moreover, our results support the validity of *FMR1* gene therapy for the treatment of FXS.

Across administration paradigms, the cellular re-expression of FMRP in AAV-*FMR1* expressing *Fmr1^-/y^* neurons remained near WT levels in most regions, consistent with prior work showing that *FMR1* promoter-based AAV expression can approximate an endogenous-like profile^48^. The inclusion of additional endogenous regulatory elements in our construct may have further supported this physiological range of expression. We acknowledge that the mild overexpression of FMRP in both WT and Fmr1-/y animals may have functional consequences that we are not aware of, however no overt toxicity or altered behavioral consequences were observed during the period examined. This is relevant in light of prior work showing that FMRP dosage is an important determinant of outcome: partial restoration can rescue phenotypes, whereas excessive overexpression can induce behavioral and molecular abnormalities^47,48,56^. Our findings are consistent with the idea that expression within an upper but permissive physiological range can support rescue, underscoring the importance of promoter choice and regulatory design in *FMR1* replacement approaches ^77^. Further work using a more titratable gene replacement strategy in addition to endogenous regulatory regions may result in even more accurately restored levels of FMRP.

This work provides new insight into how FMRP loss alters sensory-driven regulation in the IC. TRAP-seq analysis shows that sound stimulation produces a larger set of significantly altered transcripts in WT mice than in *Fmr1^-/y^* mice, indicating a more restricted stimulus-associated molecular response to sound despite the hyperactivation observed by *cfos^+^* quantification (**Fig. 3**). This pattern may reflect impaired activity-dependent regulation, however the basal changes in the translation of many of these targets suggests a state that limits further induction, consistent with prior evidence that elevated baseline protein synthesis impairs stimulus-dependent translational responses after loss of FMRP ^14,64,65,78^. In WT mice, auditory stimulation engages a coordinated program enriched for myelination, dendritic morphogenesis, vesicle transport, and oxidative phosphorylation, whereas *Fmr1^⁻/y^* mice show fewer significantly regulated transcripts, with loss of synaptic and myelination-related induction together with ectopic engagement of catecholamine transport, immune signaling, and endothelial-associated pathways. Re-expression of FMRP in AAV-*FMR1* treated *Fmr1^⁻/y^* mice restores a more WT like sound-evoked program in the IC, including genes involved in synaptic vesicle trafficking, myelination, GPCR signaling, and cytoskeletal remodeling (**Fig. 2**). Together, these findings support a model in which loss of FMRP causes a saturation of translation that prevents recruitment of activity-dependent molecular programs required for appropriate auditory circuit regulation.

Our experimental framework allows for a direct interrogation of how FMRP re-expression changes the aberrant translation of mRNAs in *Fmr1^-/y^* IC neurons. This comparison is essential for understanding what changes are, and are not, required for restoring normal activity in the *Fmr1^-/y^* auditory circuit. Comparison of the most significantly altered TRAP transcripts induced by sound in WT mice and restored by FMRP re-expression in *Fmr1^⁻/y^* mice implicated 6 candidates, among which *Npas4* was particularly notable given its established role in activity-dependent inhibitory plasticity ^67,79^. Further investigation of the TRAP-seq datasets shows that many of the downstream targets of *Npas4* are similarly insufficiently induced by stimulation in the *Fmr1^-/y^* IC, including those implicated in inhibitory synapse formation (**Fig. 2**). As TRAP is a reflection of both translation and RNA abundance, further work is needed to clarify whether the deficit in *Npas4* induction in the absence of FMRP is due to reduced translation, or an impairment in transcription or RNA stability. Our RNAscope experiments suggest that changes in *Npas4* RNA abundance are at least partially contributing to the changes seen using TRAP-seq.

The reduced recruitment of *Npas4* suggests insufficient stabilization of synaptic architecture contributing to inhibition. This would be consistent with the impaired activation of VGAT+ neurons in the *Fmr1^-/y^* IC, however future work is needed to precisely define this deficit. The IC is comprised of heterogeneous interneuron subtypes that were not resolved here ^70,80^. Distinct circuit organization and connectivity are associated with these subtypes, among the interneurons subtypes, NPY GABAergic neurons are particularly relevant as they regulate IC excitatory circuits and provide commissural inhibition ^81–83^. NPY itself is a neuropeptide associated with reduced neuronal excitability and seizure suppression, including human temporal lobe epilepsy tissue, where NPY suppress epileptiform activity ^84^. Resolving which of these subtypes is specifically impaired in the *Fmr1^-/y^* IC, and restored with FMRP re-expression, is an important next step. Moreover, identifying the regional distribution of neuronal activation within the central nucleus involved in ascending and descending information, and the dorsal and external nuclei involved in intra-IC coordination, will be important for determining the impact of *Fmr1* loss on auditory processing versus seizure generation ^30,85^.

The reduction in seizures seen with local IC injection of AAV-*FMR1* somewhat refines the circuit locus of AGS generation due to *Fmr1* loss. Following neonatal ICV delivery, FMRP is broadly re-expressed in both the IC and in ascending and descending projection areas, making it difficult to pinpoint the mechanism restoring normal activity. In contrast, the local adult delivery of AAV-*FMR1* to the IC results in a more restricted re-expression: neurons within the IC, and their ascending outputs to thalamocortical targets, can express FMRP, whereas upstream cochlear inputs and descending cortical projections to the IC are not directly corrected. Although this does not isolate the relevant synaptic pathway, it narrows the critical site of action to the IC and suggests that FMRP function within IC-centered circuitry is sufficient to substantially reduce AGS susceptibility in adulthood. These findings are also relevant to questions of reversibility and treatment timing. Neonatal ICV delivery produced robust rescue with both shorter and longer expression windows, and localized adult delivery to the IC was sufficient to reduce AGS. These observations suggest that some aspects of circuit dysfunction remain modifiable beyond early developmental stages.

Overall, our findings support a model in which FMRP preserves the dynamic range of basal and activity-dependent molecular responses in the IC. Loss of FMRP shifts this equilibrium toward a pre-engaged basal state, compresses translational and transcriptional responsiveness, weakens inhibitory recruitment, and biases the circuit toward hyperexcitability and AGS susceptibility. Re-expression of FMRP restores these processes, linking activity-dependent gene regulation to circuit stability and behavioral rescue in FXS. Given that FXS patients show sensory hypersensitivity and exaggerated auditory responses, including abnormal EEG responses to sound ^7,16–19^, the ability to modify an auditory hypersensitivity-related phenotype in adulthood has broad translational relevance.

## Acknowledgements

Special thanks to Manuela Rizzi, Wonsuk Lee and Scott Noble for help with ICV. We thank Laura Kaminioti-Dumont for help with vector and Sophie R Thomson for assistance with copy number analysis. Funding was provided by Wellcome Trust grant 219556/Z/19/Z (E.K.O.), William Hearst Fund 2026 (B.M.), and Simons Initiative for the Developing Brain (SIDB) (S.R.C.). We thank the IMPACT facility at the University of Edinburgh for imaging resources, and the Cellular Imaging Core at Boston Children’s Hospital funded by NIH IDDRC P50 HD105351.

## Author Contributions

E.K.O., B.M., and A.S. conceptualized the study and prepared the manuscript. S.R.C. and his laboratory provided the vector and contributed to the collaborative development of the study. B.M. performed ICV injections, stereotaxic injections, immunostaining, AGS, TRAP, qPCR, RNAscope, imaging and respective analysis. A.S. performed bioinformatics analyses of TRAP-seq datasets and interpretation of the study. R.H. designed the vector. J.S. and K.G. contributed to ICV injections and to the methodology of vector-related experiments. F.A. contributed to ICV injections. B.M. designed, performed, and analyzed AGS experiments with assistance from S.R.L. E.K.O., B.M. and A.S. wrote the manuscript with input from all authors.

## STAR Methods

### Animals

*Fmr1^-/y^* mice (JAX 004624) were maintained on the FVB/129 background with *Fmr1*^+/-^ females crossed with WT males. *Fmr1^-/y^* and WT TRAP mice were obtained from a F1 cross of Fmr1+/-females and Snap25-EGFP/Rpl10a males (JAX **030273).** All experiments were carried out using male littermate mice and studied with the experimenter blind to genotype. All procedures at BCH were performed in accordance with IACUC protocol 00002116. Procedures performed at University of Edinburgh were in accordance with ARRIVE guidelines and the regulations set by the University of Edinburgh and the UK Animals Act 1986. At both institutions mice were group housed (5 maximum) with unrestricted food and water access and a 12h light-dark cycle.

### Viral construct

The gene therapy vector contains human *FMR1* isoform 7 positioned between a 1050 bp fragment of human *FMR1* promoter and a 1400 bp fragment of human *FMR1* 3’UTR that includes a predominant *FMR1* polyadenylation signal. A Myc tag was included to track expression, and the construct flanked by wild-type AAV2 5’ inverted terminal repeats (ITRs). A null control vector contained a DNA stuffer sequence inserted downstream of a ubiquitous CMV promoter and SV40 polyadenylation signal. Both constructs were packaged into AAV9 capsid by Viral Vector Production Unit, Barcelona, using a Baculovirus Expression Vector System using Spodoptera frugiperda (Sf9) insect cells, and formulated in a PBS buffer containing 0.001% Poloxamer 188. Viral titer was determined using a qualified digital droplet PCR (ddPCR) assay.

### Intracerebroventricular injections

Male neonatal pups (P0-P2) were exposed to anesthesia (1.5-2% isoflurane), and bilaterally injected with AAV9-*FMR1*, AAV-NULL or PBS via the intracerebroventricular (ICV) route approximately 2mm anterior to lambda and approximately 3mm in depth, as in previous work ^19^. Virus was used at 3E11 vg/mouse.

### Stereotaxic injections

Bilateral IC injections were performed using standard aseptic techniques as in previous work^71^. P50-53 *Fmr1^-/y^*and WT mice were anesthetized with isoflurane (2-4%), and IC injection sites identified using stereotaxic coordinates (relative to the lambda suture: y: 0.9 mm; x: ± 1.0; z: 1.75). 500 nL of AAV9-*FMR1* (1.81E10 vg) or PBS was delivered per hemisphere in a rate of 100 nL/min. Post-surgical injection of analgesic (Vetergesic) and 0.9% NaCl was delivered subcutaneously. After injections were completed, the scalp was sutured with Ethilon 6–0 (0.7 metric) nylon sutures (Ethicon USA LLC), and the wound was treated with 0.5 – 1 ml 2% Lidocaine hydrochloride jelly (Akorn Inc).

### Immunostaining

Fixed brain sections (50µm) were washed with 0.3M PBS-tween(T), and antigen retrieval performed by incubating in (10 mM) sodium citrate buffer, pH 6 for 30 min at 85-95°C. Sections were then incubated in block solution (0.3M PBST with 5% normal goat serum (NGS)) for 1 h, room temperature and incubated overnight at 4°C with antibodies to FMRP (4 μg/μl, DSHB 2F5-1), NeuN (1:500, ABN91 Merck) or Myc (1:500, ab9106 Abcam) diluted in block solution. After washing 3X PBST, sections were incubated with secondary antibodies conjugated to Alexa Fluor 488 (A-11008), Alexa Fluor 568 (A-11041) and Alexa Fluor 647 (A-21236) (diluted 1:500 in block solution) for 3 h at room temperature, washed, and mounted on microscope slides using FluorSaveTM (Millipore).

Confocal imaging was performed using a Leica TCS SP8 with a Leica HC Plain Apochromat 10X/0.40 CS2 or 63X objectives. Stacks of images were acquired at 40 μm with 2.5 μm z-steps, with higher magnification images resolved with a 2X zoom and 0.5 μm z-steps. Identical imaging settings were used for all conditions within an experiment for both acquisition and analysis. Quantification was performed using Imaris 10.0.0 with the experimenter blinded to treatment and genotype. For “per neuron quantification”, a mask was applied using NeuN signal and define individual neurons, from top to bottom. FMRP and Myc Sum intensity values were normalized to NeuN volume. A minimum of 15 neurons were analyzed for each condition.

### TRAP-seq

TRAP-Seq was performed on IC from cohorts of WT and *Fmr1^-/y^*animals heterozygous for Snap25-EGFPL10a, 3 weeks after ICV injection of AAV-*FMR1* or AAV-null virus. Equal groups were either left in the home cage or exposed to a sound stimulus consisting of the same mixed-frequency sample used to evoke AGS ^63^, but limited in duration to avoid behavioral seizure manifestation. Animals were euthanized 30 min after stimulation, and IC tissue dissected and homogenized in ice-cold lysis buffer (20 mM HEPES, 5 mM MgCl2, 150 mM KCl, 0.5 mM DTT, 100 μg/ml cycloheximide, RNase inhibitors and protease inhibitors), and centrifuged at 1000×g for 10 min. Supernatants were then extracted with 1% NP-40 and 1% DHPC on ice and centrifuged at 20,000×g for 20 min. Supernatants was incubated with streptavidin/protein L-coated Dynabeads bound to anti-GFP antibodies (HtzGFP-19F7 and HtzGFP-19C8, Memorial Sloan Kettering Centre) overnight at 4 °C with gentle mixing to isolate translating ribosome-bound mRNA. Anti-GFP beads were washed with high salt buffer (20 mM HEPES, 5 mM MgCl2, 350 mM KCl, 1% NP-40, 0.5 mM DTT, and 100 μg/ml cycloheximide) and RNA was eluted from all samples using PicoPure RNA isolation kit (ThermoFisher Scientific) according to the manufacturer’s instructions. RNA with RIN > 7 was prepared for sequencing using strand-specific RNA-seq libraries for Illumina ® sequencing with the SMART- seq v4 Ultra Low Input Kit (Takara).

### RT-qPCR

RNA for each sample was converted into cDNA using Superscript VILO cDNA Synthesis Kit (Life Technologies) and RT-qPCR was performed using Quantitect SYBRgreen qPCR master mix (Qiagen) according to the manufacturer’s instructions. Samples were prepared in triplicate in 96-well reaction plates and run on a StepOne Plus (Life Technologies). Primer sequences were used as follow: Bactin: F- CAC CAC ACC TTC TAC AAT GAG; R- GTC TCA AAC ATG ATC TGG GTC, Ppib1: F- CAGCAAGTTCCATCGTGTCA; R- GATGCTCTTTCCTCCTGTGC, Gfap: F-TCC TGG AAC AGC AAA ACA AG; R- CAG CCT CAG GTT GGT TTC AT, Snap25: F- TGA GGA AGG GAT GGA CCA AA; R- CCT GAT TAT TGC CCC AGG CT, Myc: F- GAT CGC TTC AGA TCA GAG TTG; R- CCT CTT CTG AGA TGA GTT TTT GTT C. Fold change was calculated using the delta-delta Ct method where the first delta was calculated as the average of b-actin and Ppib1, and the second delta was calculated as the average of all samples run together for each experiment to avoid canceling out the variation in control samples. Values were then normalized to the mean of all control samples for graphical purposes. For determination of Myc copy number, a set of 6 standards were prepared in LE-TE buffer by 10-fold dilution, and Myc transcript number extrapolated from the standard curve.

### RNA-Seq library preparation and analysis

Low-input mRNA libraries were prepared by Psomagen using the Takara SMART-Seq mRNA LP kit and sequenced on an Illumina NovaSeq X Plus platform to generate 150 bp paired-end reads. Raw reads were processed with Trim Galore v0.6.6 ^86^ to remove Illumina adapter sequences and low-quality bases using the parameters -q 30 --length 50 --illumina --paired --trim-n. Transcript abundance was quantified using Salmon v1.10.1 ^87^ for filtered reads against a transcriptome index in quasi-mapping mode with automatic library type detection (-l A) and sequence- and GC-bias correction enabled (--validateMappings, --seqBias, --gcBias). Salmon quantification outputs were then used for downstream gene-level expression analysis.

### Differential expression analysis on TRAP-Seq data

Transcript-level abundance estimates were imported into DESeq2 (v1.46.0) using tximport, with transcript counts summarized to the gene level using a custom transcript-to-gene annotation table. Metadata for each sample, including biological replicates, condition, stimulation state, batch and treatment were incorporated into the DESeq2 analysis framework. A combined group variable was created from condition, stimulation, and treatment, and differential expression analysis was performed using DESeq2 with the design formula ∼ batch + group. Genes with low expression were filtered prior to analysis by requiring a minimum of 10 counts in at least half of the samples. Size factors and dispersions were estimated using DESeq2, followed by negative binomial generalized linear modeling and Wald significance testing.

### Gene set Enrichment and Gene Ontology analysis

GSEA v4.3.2 Mac App was downloaded from website (https://www.gsea-msigdb.org/gsea/) and annotated list of ontology gene sets (m5.go.v2025.1) for biological, molecular and cellular pathways were used from Molecular Signature Database - MSigDB (v2025.1.Mm). GSEA analysis was performed using GSEAPreranked method, where genes were ranked by fold change and a ‘classic’ enrichment statistic was used to remove the magnitude bias of ranking metric. Minimum of 20 and maximum of 500 was defined as cutoff for number of genes in a gene set identification with maximum 1000 permutations. GSEA comparison across datasets was performed for significant terms using a cutoff of Nominal P values (P < 0.01) or FDR (Padj < 0.1) as indicated and ranked by normalized enrichment score (NES). Tables summarizing these GSEA categories are supplied as extended data. Gene ontology analysis was performed using ClusterProfileR v4.14.6 ^88^ on significantly (P < 0.01) up- or downregulated genes with a background of all identified genes in respective dataset.

### Correlation and slope comparison analysis

To compare the relationship between translational responses across conditions, genes detected in both comparisons were plotted in a scatter plot by their corresponding log2 fold-change value. Each point represents one gene on the scatterplot, pairwise correlations were calculated from the indicated DESeq2 comparisons. Pearson correlation coefficients were used to quantify the strength and direction of the linear relationship between comparisons, unless otherwise stated. Linear regression lines were fitted to visualize the overall relationship between the two response profiles.

To statistically compare whether the relationship between two datasets differed across conditions, we used a generalized linear model with an interaction term. For each analysis, the two y-axis datasets were combined into a single dataframe, using the same x-axis variable for both groups and a categorical variable indicating the comparison group. The model was specified as: glm(y ∼ x * comparison, data = plotdf, family = gaussian()). In this model, the x:comparison interaction term tests whether the two regression slopes are statistically different.

### RNAscope

Frozen sections (16 µm) from WT and *Fmr1^-/y^* mice were cut coronally in cryostat and mounted onto Superfrost Plus microscope slides. Sections were fixed in 4% PFA for 15 minutes, dehydrated in increasing concentrations of ethanol for 5 minutes, air-dried, and then pretreated with protease for 30 minutes at room temperature. For RNA detection, sections were incubated in different amplifier solutions in a HybEZ hybridization oven (Advanced Cell Diagnostics) at 40°C. Synthetic Oligonucleotides complementary to the sequence range of 793-1795 (Mm-*Npas4*-C2), and range 407-1447 for (Mm-*cfos*-C2), range 1986-2998 for (Mm-*VGLUT2*-C2), and range 894-2037 for (Mm-*VGAT*-C2) (Advanced Cell Diagnostics) were used as probes. The labeled probes were conjugated to opal Dye 520, 570 and 620, after which labeled probe mixtures were hybridized by incubating with slide-mounted sections for 2 hours at 40C. Nonspecifically hybridized probes were removed by washing the sections three times for 2 minutes each with 1X wash buffer at room temperature, followed by incubation with Amplifier 1-FL for 30 minutes, Amplifier 2–FL for 15 minutes, Amplifier 3–FL for 30minutes, and Amplifier 4 AltB–FL for 15minutes at 40°C. Each amplifier was removed by washing with 1X wash buffer for 2 minutes at room temperature. The slides were imaged using a confocal microscope (Leica sp8) with a 10-x objective lens; all image settings were kept constant.

### Audiogenic seizures

Experiments were performed as previously described ^63,71^. Naive male FVB/N P19–60 or Snap25-EGFPx FVB/N mice were transferred to a quiet (<60 dB ambient sound) room for 1 h. For testing, animals were moved to a transparent test chamber (39×3×23 cm) equipped with speakers and a camera and allowed to habituate for 1 min. Audiogenic stimulation (recorded sampling of a modified personal alarm) was passed through an amplifier and 2x 50 W speakers (KRK Rokit RP5 G3 Active Studio Monitor) to produce a stimulus of 120 dB for a maximum of 2 min. During each testing session mice from both genotype and treatment groups were included, with experimenter blind to genotype and treatment. A decibel meter was placed on a clear plexiglass sheet 28 cm above the arena, in order to monitor sound output at a fixed distance throughout the experiment. Incidence and severity of seizures was scored. AGS severity was calculated as follows: wild running (WR; pronounced, undirected running and thrashing), clonic seizure (violent spasms accompanied by loss of balance), or tonic seizure (postural rigidity in limbs). Any animal that reached tonic seizure was immediately humanely euthanized. Latency was measured as the number of seconds between onset of the AGS stimulus and appearance of the first seizure.

### Data Analysis and Quantification

All statistical analyses were performed using GraphPad Prism or R. AGS incidence scores were analyzed using Fisher’s Exact test. For RNAscope experiments significant effects were determined using repeated-measures 2-way ANOVA or 3-way ANOVA followed by post-hoc tests. For RNA-seq datasets, differential expression was determined using DESeq2 using the default cutoff for significance (adjusted P value < 0.1). For GO and GSEA, significance was determined by nominal P value, and adjusted p-values with FDR cutoff where indicated. For overlap analyses, significant features from the two datasets being compared were converted into binary membership vectors within a shared background universe of size N, where features represented genes, GO terms, or GSEA terms depending on the analysis. For two feature sets A and B, with |A|=a, |B|=b, and observed overlap |A∩B|=k overlap significance was estimated using a one-sided hypergeometric test:

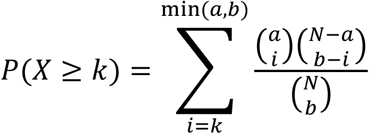

## Supplemental Information

**Figure S1. Related to Fig. 1.**
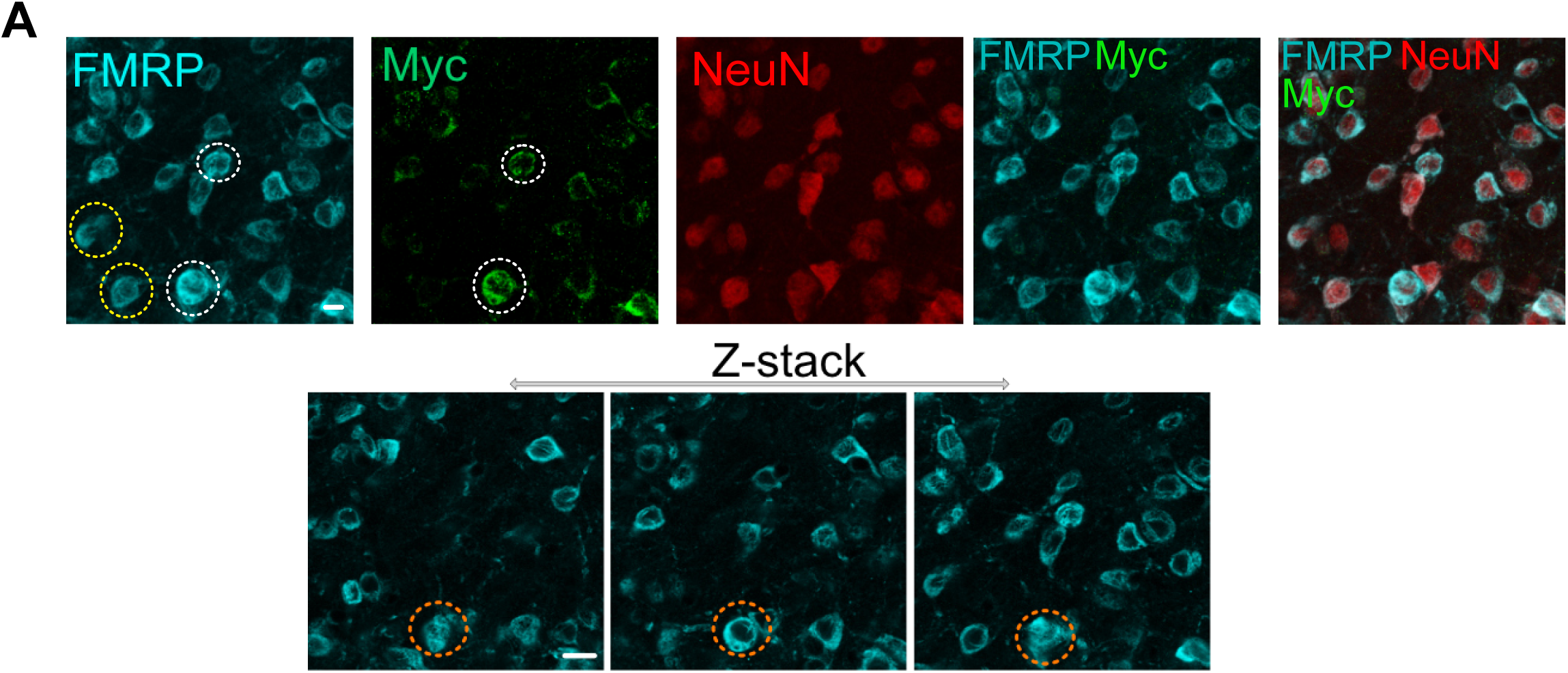
FMRP per neuron was quantified in acquired z-stacks as mean intensity in NeuN+ cells. Scale bar = 10 µm. Yellow circles highlight FMRP-positive/Myc-negative neurons, whereas white circles indicate Myc-positive neurons. Orange circles illustrate the identification of neurons across the z-stack.

**Figure S2. Related to Fig. 2.**
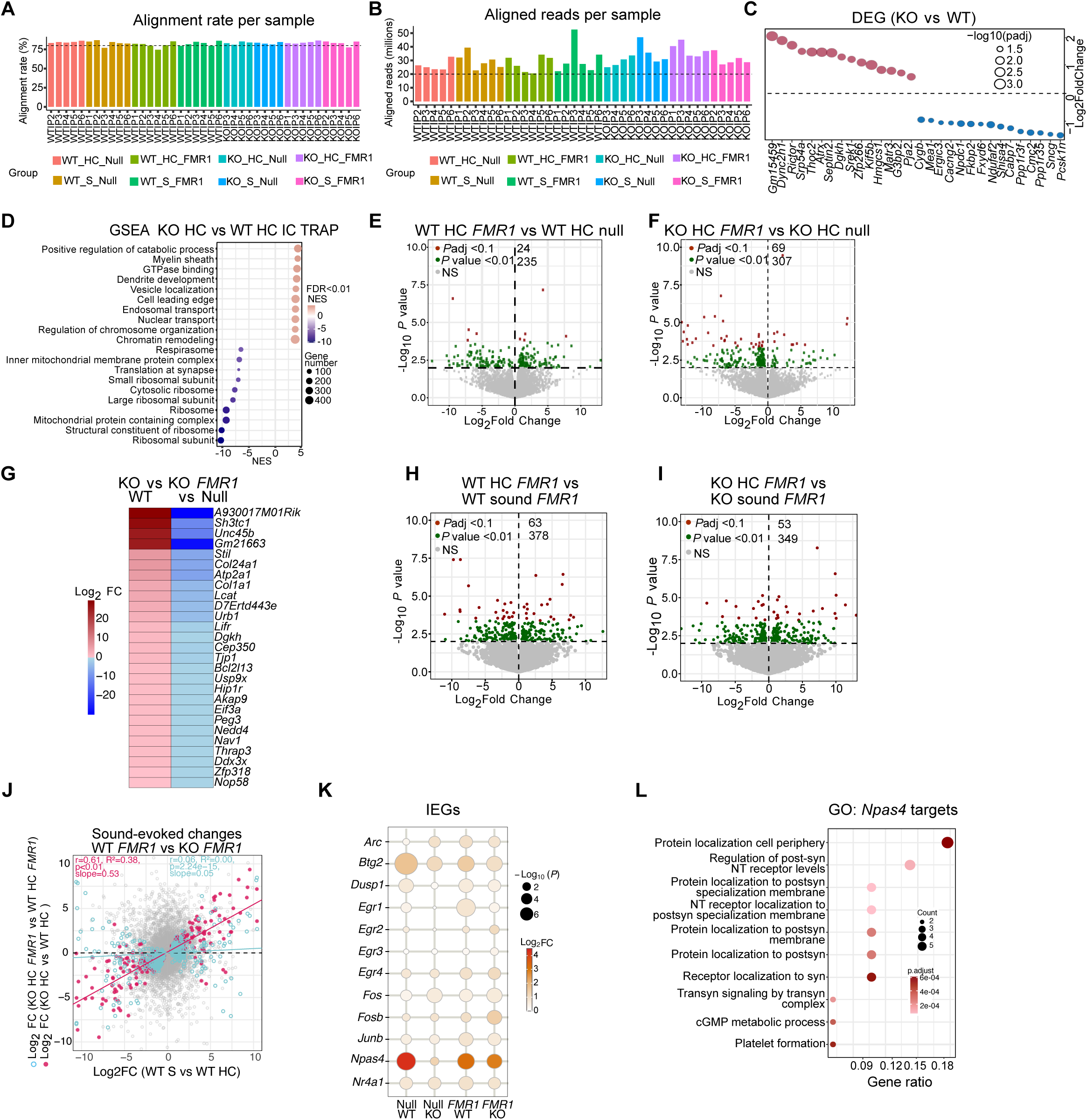
**(A)** Alignment rate across IC TRAP-seq samples grouped by genotype, stimulation condition, and viral treatment. Dashed line indicates 80% alignment rate as a visual reference. **(B)** Number of aligned reads per sample, shown in millions, across the same experimental groups. Dashed line indicates 20 million aligned reads as a visual reference. **(C)** Ranked fold-change plot showing the most significant (Padj < 0.1) translational changes in KO HC relative to WT HC. (**D**) Gene set enrichment analysis of differentially expressed genes in KO HC compared with WT HC IC TRAP samples, highlighting positively and negatively enriched biological processes/pathways. Dot color represents normalized enrichment score (NES), and dot size represents the number of genes contributing to each term. Terms shown pass the indicated FDR threshold. **(E)** Volcano plot showing differential gene expression in WT HC samples receiving *FMR1* compared with WT HC Null-control samples. **(F)** Volcano plot showing differential gene expression in KO HC samples receiving *FMR1* compared with KO HC Null-control samples. (**G**) Heatmap of basal changes in KO HC versus WT HC that are reversed by *FMR1* re-expression, highlighting synaptic, transporter, protein-binding and vesicle associated functions. **(H)** Volcano plot showing differential gene expression in WT HC samples receiving *FMR1* compared with WT sound receiving *FMR1*. **(I)** Volcano plot showing differential gene expression in KO HC samples receiving *FMR1* compared with KO Sound receiving *FMR1* samples. Genes are colored by significance category based on Padj < 0.1 or nominal P value < 0.01. (J) Scatter plot comparing sound-evoked changes in WT to basal changes in KO *FMR1* reveals no correlation (r = 0.06, R^2^ = 0), whereas a positive correlation is seen with KO NULL (r = 0.61, R^2^ = 0.38, *p < 0.01). (**K**) Dot plot of immediate early gene responses across sound stimulated samples. Dot size indicates significance and color indicates log2 fold change. (**L**) Gene Ontology analysis of known *Npas4* target genes upregulated by sound in WT mice.

**Figure S3. Related to Fig. 3.**
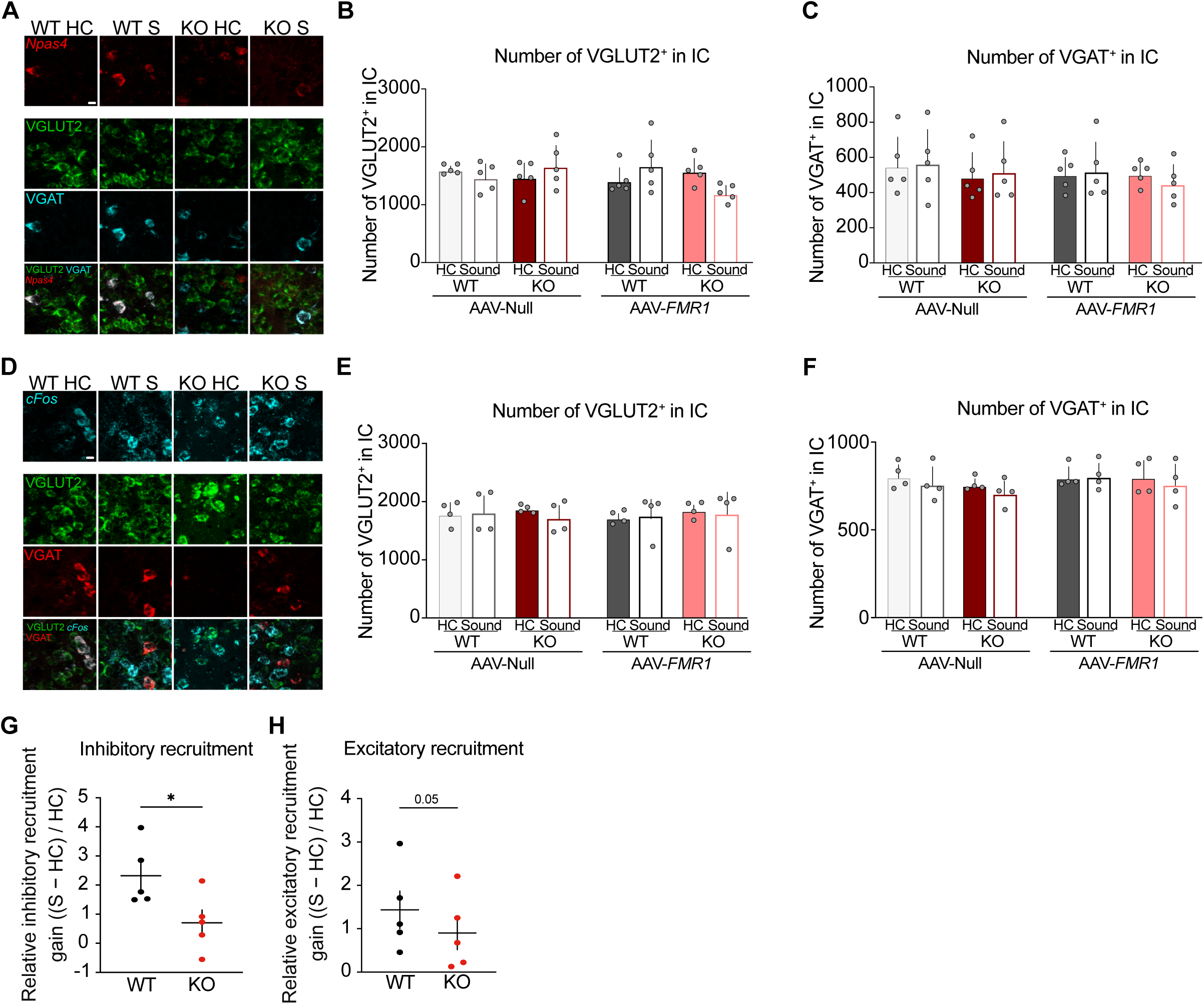
(**A**) Representative 63× confocal images of *Npas4*, VGLUT2, and VGAT. Scale bar = 10 µm. (**B**) RNAscope for *Npas4*: Number of VGLUT2^+^ in IC across conditions. (**C**) RNAscope for *Npas4*: Number of VGAT^+^ in IC across condition. n = 5 littermate pairs. Bars indicate mean ± SEM. (**D**) Representative 63× confocal images of *cfos*, VGLUT2 and VGAT. Scale bar=10um. (**e**) RNAscope for *cfos:* Number of VGLUT2^+^ in IC across conditions. (**F**) RNAscope for *cfos*: Number of VGAT^+^ in IC across condition. n = 5 littermate pairs. Bars indicate mean ± SEM. (**G**) Normalized change in VGAT⁺ inhibitory and VGLUT2⁺ excitatory (**H**) recruitment, calculated (S − HC) / HC. Student paired t-test, * p = 0.0294 for inhibitory recruitment and excitatory recruitment p = 0.0584. Each dot represents one mouse; bars indicate mean ± SEM.

**Table S1:** DESeq2 results for all transcripts in TRAP-seq across the comparison groups.

**Table S2:** Gene ontology analysis terms for WT HC vs KOHC differentially expressed genes.

**Table S3:** GSEA analysis for IC TRAP-seq comparison groups.

**Table S4:** Gene ontology analysis terms for *Npas4* targets.

**Table S5**: Number of mice tested and the AGS severity distribution across the three paradigms.

**Video S1. P60 WT PBS.** Shows 1 min habituation followed by 2 min AGS stimulation of a P60 WT mouse injected with PBS.

**Video S2. P60 WT AAV-NULL.** Shows 1 min habituation followed by 2 min AGS stimulation of a P60 WT mouse injected with AAV-NULL.

**Video S3. P60 WT AAV-*FMR1*.** Shows 1 min habituation followed by 2 min AGS stimulation of a P60 WT mouse injected with AAV-*FMR1*.

**Video S4. P60 *Fmr1^-/y^* PBS.** Shows 1 min habituation followed by 2 min AGS stimulation of a P60 *Fmr1^-/y^* mouse injected with PBS.

**Video S5. P60 *Fmr1^-/y^* AAV-NULL.** Shows 1 min habituation followed by 2 min AGS stimulation of a P60 *Fmr1^-/y^* mouse injected with AAV-NULL

**Video S6. P60 *Fmr1^-/y^* AAV-*FMR1*.** Shows 1 min habituation followed by 2 min AGS stimulation of a P60 *Fmr1^-/y^* mouse injected with AAV-*FMR1*.

**Video S7. P60 WT PBS, IC injection.** Shows 1 min habituation followed by 2 min AGS stimulation of a P60 WT mouse injected with PBS in IC.

**Video S8. P60 WT AAV-*FMR1*, IC injection.** Shows 1 min habituation followed by 2 min AGS stimulation of a P60 WT mouse injected with AAV-*FMR1* in IC.

**Video S9. P60 *Fmr1^-/y^* PBS, IC injection.** Shows 1 min habituation followed by 2 min AGS stimulation of a P60 *Fmr1^-/y^* mouse injected with PBS in IC.

**Video S10. P60 *Fmr1^-/y^* AAV-*FMR1*, IC injection.** Shows 1 min habituation followed by 2 min AGS stimulation of a P60 *Fmr1^-/y^*mouse injected with AAV-*FMR1* in IC.

**Video S11. P21 WT PBS.** Shows 1 min habituation followed by 2 min AGS stimulation of a P21 WT mouse injected with PBS.

**Video S12. P21 WT AAV-NULL.** Shows 1 min habituation followed by 2 min AGS stimulation of a P21 WT mouse injected with AAV-NULL.

**Video S13. P21 WT AAV-*FMR1*.** Shows 1 min habituation followed by 2 min AGS stimulation of a P21 WT mouse injected with AAV-*FMR1*.

**Video S14. P21 *Fmr1^-/y^* PBS.** Shows 1 min habituation followed by 2 min AGS stimulation of a P21 *Fmr1^-/y^* mouse injected with PBS.

**Video S15. P21 *Fmr1^-/y^* AAV-NULL.** Shows 1 min habituation followed by 2 min AGS stimulation of a P21 *Fmr1^-/y^*mouse injected with AAV-NULL.

**Video S16. P21 *Fmr1^-/y^* AAV-*FMR1*.** Shows 1 min habituation followed by 2 min AGS stimulation of a P21 *Fmr1^-/y^*mouse injected with AAV-*FMR1*.

